# tRNA hypomodification facilitates 5-fluorocytosine resistance via cross-pathway control system activation in *Aspergillus fumigatus*

**DOI:** 10.1101/2024.02.19.578369

**Authors:** Alexander Bruch, Valentina Lazarova, Maximilian Berg, Thomas Krüger, Sascha Schäuble, Abdulrahman A. Kelani, Birte Mertens, Pamela Lehenberger, Olaf Kniemeyer, Stefanie Kaiser, Gianni Panagiotou, Fabio Gsaller, Matthew G. Blango

## Abstract

Increasing antifungal drug resistance is a major concern associated with human fungal pathogens like *Aspergillus fumigatus*. Genetic mutation and epimutation mechanisms clearly drive resistance, yet the epitranscriptome remains relatively untested. Here, deletion of the *A. fumigatus* tRNA-modifying isopentenyl transferase ortholog, Mod5, led to altered stress response and unexpected resistance against the antifungal drug 5-fluorocytosine (5-FC). After confirming the canonical isopentenylation activity of Mod5 by LC-MS/MS and Nano-tRNAseq, we performed simultaneous profiling of transcriptomes and proteomes to reveal a comparable overall response to 5-FC stress; however, a premature activation of cross-pathway control (CPC) genes in the knockout was further increased after antifungal treatment. We identified several orthologues of the *A. nidulans* Major Facilitator Superfamily (MFS) transporter *nmeA* as specific CPC-client genes in *A. fumigatus*. Overexpression of Mod5-target tRNA^Tyr^_GΨA_ in the Δ*mod5* strain rescued select phenotypes but failed to reverse 5-FC resistance, whereas deletion of *nmeA* largely, but incompletely, reverted the resistance phenotype, implying additional relevant exporters. In conclusion, 5-FC resistance in the absence of Mod5 and i^6^A likely originates from multifaceted transcriptional and translational changes that skew the fungus towards premature CPC-dependent activation of antifungal toxic-intermediate exporter *nmeA*, offering a potential mechanism reliant on RNA modification to facilitate transient antifungal resistance.

## INTRODUCTION

Severe fungal infections have increased constantly in recent decades and represent a substantial threat to human health, resulting in over a billion infections and roughly 3.75 million deaths per year (GAFFI.org, (1–3)). In 2022, this danger was recognized by the World Health Organization (WHO) as a prioritized list of fungal species, based on infectivity, severity of infection, and propensity for antifungal resistance (4). Among the critical priority group of fungal pathogens is *Aspergillus fumigatus*, a filamentous saprophytic fungus that resides in the environment but is able to reach the lungs of human hosts as airborne asexually produced spores, termed conidia (5). In healthy individuals, the innate immune system eliminates invading spores, whereas heavily immunocompromised patients are susceptible to outgrowth of the conidia and dangerous invasive infections (6,7). Treatment of these infections relies on a limited set of antimycotics, including the echinocandins, polyenes, and azoles, of which the latter represents the first-line medication used to treat *A. fumigatus* infections (8,9). Other potential drugs like 5-fluorocytosine (5-FC; flucytosine) are normally used in combinatorial antifungal treatments but are not used to treat *A. fumigatus* infections due to swift development of resistance (5,8,10). Additionally, development of antifungal resistance is further amplified by heavy utilization of molecularly similar drugs in agriculture, creating a selective environmental pressure (11,12). Clearly, a better understanding of the molecular mechanisms underlying resistance and virulence are needed to combat the problem of human fungal infections.

Many studies have investigated genomic-encoded resistance, but much less is known about the role of the more than 170 known RNA chemical modifications, termed the epitranscriptome, in control of resistance and virulence (13,14). RNA modifications are particularly prevalent on transfer RNAs (tRNA), where these range from simple methylations (e.g., 5-methyl cytosine=m^5^C) or isomerisations (e.g., pseudouridine=Ψ) to complex chemical groups like N^6^- isopentenyl adenosine (i^6^A). Modifications fulfil a variety of functions, for example by supporting the integrity of the tRNA tertiary structure or translational fidelity (13,15). The highly-conserved i^6^A modification is installed at position 37 of select adenosines in the anticodon stem-loop by isopentenyl transferases (IPTase) to support base stacking, prevent cross- anticodon stem-loop base pairing between U33 and A37, and fine-tune codon-anticodon interactions (16–20).

The i^6^A modification has been investigated in multiple pathogenic model systems. Deletion of the IPTase *miaA* in the bacterium *Shigella flexneri* resulted in the absence of i^6^A37 and its derivative ms^2^i^6^A37 and strongly decreased the expression of different virulence factors (21). Interestingly, these changes only occurred on the translational but not transcriptional level, indicating a regulatory dependency of pathogenicity by i^6^A/ms^2^i^6^A-modified tRNAs. Recently, the role of MiaA and corresponding tRNA modification in stress adaptation and virulence was shown for extraintestinal pathogenic *Escherichia coli* (ExPEC) (22). Oscillation of the modifier during stress situations clearly indicated an active adaptation of the i^6^A/ms^2^i^6^A tRNA modification pattern in ExPEC that affected the translational capacity of the bacteria and modulated the composition of the proteome. This was even more apparent when the MiaA level was artificially changed by either ablation or overexpression (22), suggesting that MiaA could function as a molecular rheostat for control of stress response.

We hypothesized that similar regulation likely exists in fungal pathogens, and a few studies have indeed reported that the epitranscriptome contributes to fungal virulence and stress resistance. For instance, a study focusing on the kinome of *Cryptococcus neoformans* revealed that specific tRNA modification-relevant kinases, namely *BUD32*, *TRM7,* and *TRM4*, are important for its pathogenicity (23). These findings were underscored by a recent report focusing on Sua5, which in cooperation with Bud32 (and other proteins), is responsible for t^6^A_37_ tRNA modification, pointing out its significance for fungal virulence (24). Similarly, several papers recently described how the modulation or ablation of mcm^5^/s^2^U34 affected the metabolism, stress adaptivity, and virulence of *Candida albicans*, *Magnaporthe oryzae,* and *A. fumigatus* (25–27). These studies suggest that anticodon stem-loop modifications, specifically at position 34 and 37, have been underestimated for their relevance in host-fungal interactions.

Little is known of the function of i^6^A37 in pathogenic fungi, with many insights of fungal IPTase- dependent modifications coming from non-pathogenic model systems like *Saccharomyces. cerevisiae* or *Schizosaccharomyces pombe* (14,16,28). In bakers’ yeast, Mod5 can localize to different cellular compartments to install i^6^A on cytoplasmic and mitochondrial tRNAs, but it also translocates into the nucleus to perform tRNA gene-mediated silencing (TGM), a secondary transcription regulatory function (29–36). Interestingly, the yeast protein also contains a prion domain that enables it to form amyloid structures (37). The shift from soluble protein to prion form (and back) was found to not only influence the modification function of Mod5 but also represents a molecular switch to adapt to azole drugs, providing a connection between tRNA modification and antifungal drug resistance (37). Despite its many functions, deletion of *MOD5* is well compensated by bakerś yeast and led to few phenotypes beyond slight sporulation defects and decreased suppressor efficiency of tRNA^Tyr^UAA (16,19). Similar to *S. cerevisiae*, deletion of the IPTase *TIT1* in *S. pombe* resulted in few phenotypes, limited to increased sensitivity against rapamycin treatment, reduced translation, and mitochondrial dysfunction when grown on non-fermentable carbon sources (38,39). Interestingly, despite multiple cytoplasmic and mitochondrial tRNAs being modified by Tit1, overexpression of the hypomodified cy-tRNA^Tyr^GΨA reverted the observed phenotypes, indicating that this tRNA was strongly affected by lack of i^6^A37 in *S. pombe* (38). In summary, proper tRNA modification is a prerequisite for pathogenesis, with surprising implications on antifungal drug resistance.

Here, we present evidence that deletion of *mod5* impacts the transcriptome, translational capacity, and stress response of the opportunistic fungal pathogen *A. fumigatus*. Surprisingly, deletion of this tRNA modification enzyme resulted in increased resistance of the fungus against the antimycotic agent 5-FC, which relied on premature activation of cross-pathway control (CPC) to drive expression of the toxic nucleobase exporter *nmeA* and facilitate antifungal drug resistance.

## MATERIAL AND METHODS

### Culture and growth conditions

*Aspergillus fumigatus* strains used in this study (**Table S1**) were cultivated on AMM agar plates according to (40) at 37°C. Spores from respective agar plates were harvested after 5 days utilizing a T-cell scraper, suspended in 10 mL sterile water, and filtered through a 30- µm cell strainer (MACS, Miltenyi Biotec GmbH; Bergisch Gladbach, Germany) to remove remaining mycelium. Spore concentration was determined using a Roche Innovatis CASY Cell Counter & Analyzer System Modell TT. For liquid cultures, 1x10^8^ or 5x10^7^ *A. fumigatus* spores were cultured in 50 mL AMM for 24 or 48 h at 37°C and 200 rpm. Mycelium was harvested using Miracloth (EMD Millipore Corp., USA) and rinsed with ultrapure Milli-Q® water (Merck Millipore, Germany). Positive *A. fumigatus* transformants were selected from AMM agar plates (containing 1 M sorbitol) that were supplemented with either pyrithiamine (0.1 µg/mL), phleomycin (80 µg/mL), or hygromycin B (200 µg/mL). *E. coli* cultures were grown in LB media at 37°C and supplemented with 60 µg/mL carbenicillin as necessary.

### Droplet stress assays

The droplet stress assays were conducted with freshly harvested spores from five-day cultures. Spore suspensions were adjusted to a concentration of 2x10^7^ spores/mL in sterile water corresponding to 1x10⁵ spores in 5 µL. Ten-fold dilutions of 10⁵-10² spores/5 µL were spotted on square petri dishes containing 50 mL of AMM, which was supplemented with 500- 750 ng/mL rapamycin, 0.3 mM 3-Amino-1,2,4 Triazole (3-AT), 0.4 µg/mL 5-fluoroorotic acid (5-FOA), 1-80 ng/µL 5-FC, 40 µg/mL hygromycin, 0.1 µg/mL itraconazole, 0.5 µg/mL tebuconazole, 0.25 µg/mL voriconazole, and 0.5-8 µg/mL caspofungin. Assays were incubated at 37°C for 48 and 72 h and documented using a Nikon D6500 AF-S NIKKOR 18-105mm f/3.5- 5 ED VR camera.

### Minimum inhibitory concentration (MIC) assays

MICs were determined by suspending wild-type and Δ*mod5* spores in 96-well plates containing RPMI 1640 Medium (Thermo Fisher Scientific), adjusted to a final concentration of 2% glucose (v/v) and grown for 48 h. Each well contained increasing concentrations of the antifungal compound, as well as positive and sterile controls up to a total volume of 200 µL. Spore dilutions of 2x10^7^/mL for both wild type and knockout were prepared in 1x RPMI 2% glucose for a final inoculum of 1x10^5^ total spores per well delivered in 5 µL. The antifungal agent 5-FC was sterilized using 0.22-µm sterile filter (Carl Roth, Germany) and diluted in 1x RPMI 2% glucose (v/v) to attain the desired concentrations. Visual assessment of hyphal growth to elucidate the MIC was performed by light microscopy.

### Strategies for genetic manipulation of *A. fumigatus*

Generation of *mod5* knockout*, AFUB_005530 (nmeA)* knockout, tRNA^Tyr^GΨA-overexpression, and *mod5* complementation constructs relied on the Gibson cloning method. The deletion approach was conducted according to (40) amplifying a selectable marker gene together with flanking sequences of about 1 kb for the *mod5* (*AFUB_093210*) or *nmeA* (*AFUB_005530*) genes, employing the primer pairs given in **Table S2**. The tRNA^Tyr^GΨA-overexpression construct was generated by cloning 3 PCR products: tRNA^Tyr^GΨA gene (*AFUB_013600*), a vector backbone (consisting of *pyrG* flanking regions; phleomycin resistance marker; and tetracycline-inducible promoter), and synthesized *Aspergillus nidulans* Ttef transcriptional terminator. tRNA^Tyr^GΨA gene and Ttef were amplified using primer pairs (tRNA^Tyr^GΨA-OE-o1 & tRNA^Tyr^GΨA-OE-o2) and (tRNA^Tyr^GΨA-OE-o3 & tRNA^Tyr^GΨA-OE-o4), respectively. The vector backbone was amplified from a previously described plasmid pSilent-*pksP* 2 (40) using primer pair tRNA^Tyr^GΨA-OE-o5 and tRNA^Tyr^GΨA-OE-o8. For the generation of the Δ*mod5* complemented strain, the *mod5* gene was PCR amplified and cloned into a pUC18 vector backbone with flanks complementary to *pyrG* and a phleomycin resistance marker (**Table S2**). Insertion of the tRNA^Tyr^GΨA-overexpression construct, *mod5*-complementation construct or deletion of *mod5* or *nmeA* in *A. fumigatus* CEA17 Δ*akuB^KU80^* was achieved by protoplast transformation and homologous recombination following the instructions of (41).

In order to generate a tuneable *nmeA* strain, an expression cassette was integrated, comprising *nmeA* coding sequence under control of the xylose-inducible promoter *PxylP* (42), at the recently established counter-selectable marker locus *cntA* (43). The respective expression cassette was amplified from the plasmid pΔfcyB_PxylP-nmeA using primers NmeA-OE-o1 and NmeA-OE-o2 (**Table S2**), adding 50 bp overhangs of 5’ and 3’ *cntA* flanks for CRISPR/Cas9-assisted–microhomology-mediated replacement of the *cntA* coding sequence. pΔfcyB_PxylP-nmeA was generated as follows. A plasmid backbone containing *PxylP* was generated by PCR from pΔfcyB_mKate2xyl using primers NmeA-OE-o3 and NmeA- OE-o4 as recently described (44). Next, *nmeA* coding sequence with 20 bp overlapping ends to the backbone was amplified from wild-type genomic DNA using primers NmeA-OE-o5 and NmeA-OE-o6 (**Table S2**) and assembled with the backbone using NEBuilder® (NEB).

1 µg of DNA were transformed into wild-type protoplasts together with a ribonucleoprotein complex comprising Cas9 (NEB) and single guide RNA (IDT) (sgRNA; 5‘- GGCGGACGACAAGGCUAUGG-3‘) following an approach described recently (45). Transformants were selected on *Aspergillus* minimal medium (AMM; (46)) supplemented with 0.1 M citric acid buffer pH 5, 1 % glucose, 20 mM ammonium tartrate, 1 M sucrose and 50 µg/mL clorgyline (Sigma) and 50 µg/mL 5-fluorouridine (TCI) for selection.

### RNA extraction and processing

For RNA extraction from fungal mycelia, 1x10^8^ spores were inoculated in 50 mL AMM and grown for 16 to 24 h (+ 4-8 h with appropriate 3-AT or 5-FC stressors, respectively) at 37°C and 200 rpm. Afterwards, mycelium was harvested using Miracloth (EMD Millipore Corp., USA), rinsed with sterile water and equal amounts of mycelium were transferred to 2-mL screw-cap tubes for RNA isolation. All centrifugation steps were performed at 4°C and full speed (20,817 x *g*). 800 µL of TRIzol (Thermo Fischer Scientific, Dreieich) were added to each tube, along with glass beads (1/3 of the tube) and the mycelium was mechanically disrupted by 3 rounds of 30 sec, 4.0 m/sec homogenization using the FastPrep-24™ Homogenizer. Subsequently, the samples were incubated for 5 min on ice, followed by an additional 5 min at room temperature. After addition of 160 µL of chloroform, the samples were vigorously vortexed for 10 sec and centrifuged for 5 min. The resulting aqueous phase containing the RNA was carefully transferred to a new RNase-free tube without disturbing the interphase. The RNA phase was then purified using an equal amount of phenol/chloroform/isoamyl alcohol and centrifuged for 5 min. The purification step was repeated 2-3 times until the interphase remained clear. RNA was further purified with 400 µL of chloroform and centrifugation for 5 min. The aqueous phase was again transferred to a new RNase-free 2-mL tube, precipitated by addition of 500 µL isopropanol for 20 min, and RNA was pelleted by 20 min centrifugation. The RNA pellet was washed with 70% ethanol (v/v) and air-dried at 37°C. Finally, RNA was dissolved in 30-50 µL DEPC water at 65°C. The RNA was purified using the RNA Clean and Concentrator™ kit (Zymo Research, USA) following the manufacturer’s instructions. Up to 10 µg of total RNA was afterwards treated with 2 U of DNase I endonuclease (Thermo Fisher) for 30-60 min at 37°C in a volume of 20 µL. RNA concentration was quantified with the Qubit^TM^ RNA BR Assay kit (Thermo Fisher Scientific) on the Qubit Flex Fluorometer (Thermo Fisher Scientific). Quality was assessed either by agarose gel electrophoresis or QUBIT^TM^ Integrity and Quality (IQ) kit (Thermo Fisher).

### tRNA enrichment from total RNA

Isolation of tRNAs from total RNA samples for the quantitative and qualitative analysis of tRNA modifications (i^6^A, Ψ, m^5^U, 5-FU) was conducted according to (47) with minor adjustments. Briefly, for tRNA isolation from *A. fumigatus* strains CEA17 Δ*akuB^KU80^* and Δ*mod5*, 1x10^8^ spores were cultured for 24 h in 50 mL of AMM at 37°C, 200 rpm. Mycelium was harvested using Miracloth (EMD Millipore Corp., USA) and crushed with mortar and pestle under constant treatment with liquid nitrogen. Approximately 0.8 mL of mycelium was transferred into a 2-mL Safe-Lock tube, mixed with an equal volume TRIzol, and snap frozen in liquid nitrogen. The samples were left on ice to thaw, mixed again, and incubated for 5 min first on ice and afterwards at room temperature. After addition of 0.2 volumes of chloroform per volume of TRIzol, the samples were vortexed and centrifuged at full speed for 5 min at 4°C. The upper aqueous phase containing RNA (ca. 600 µL) was transferred into a new tube and centrifuged again (15,000 rpm, 30 min, 4°C) resulting in a supernatant that was collected in a new tube (ca. 500 µL). For subsequent total tRNA precipitation, 3/4 vol. 8 M LiCl was added to the samples and incubated for 2 h (or longer) at -20°C. Samples were centrifuged (14,000 rpm, 15 min, 4°C) to separate larger RNA molecules and the supernatant containing small RNAs was transferred into a new tube. The precipitation was repeated once to improve purification. The supernatant was mixed with 1/10 volume ammonium acetate, pH 7.0 and 2.5 volumes of ice-cold 100% EtOH p.a. to be incubated overnight at -20°C. Precipitated tRNA was collected by centrifugation (14,000 rpm, 30 min, 4°C). The pellet was washed twice briefly with 75% (v/v) EtOH and dry spun to remove residual ethanol. The dried RNA was resuspended in 50 µL DEPC-H2O. The tRNA samples were quantified with Qubit^TM^ RNA BR Assay kit (Thermo Fisher Scientific) on the Qubit Flex Fluorometer (Thermo Fisher Scientific) and stored at -80°C.

### tRNA^Tyr^GUA isolation for nucleoside LC-MS/MS

tRNA^Tyr^GUA purification was performed according to published procedures (48,49). 1 µg of total tRNA was mixed with 100 pmol reverse complementary, biotinylated DNA oligonucleotide (Sigma Aldrich, St. Louis, MO, USA; sequence: [Btn]CCCGAGCCGGAATCGAGCCG, [Btn] = Biotin-Tag) in a total volume of 100 µL 5X SSC buffer (0.75 M NaCl, 75 mM trisodium citrate pH 7). The mixture was incubated for 3 min at 90°C for denaturation, followed by a hybridization step for 10 min at 65°C. For each sample, 25 µL Magnetic Dynabeads® Myone^TM^ Streptavidin T1 (Thermo Fisher Scientific, Waltham, MA, USA) were primed three times using Bind and Wash buffer (5 mM Tris-HCl pH 7.5, 0.5 M EDTA, 1 M NaCl) and once using 5X SSC buffer. An aliquot of 25 µL magnetic beads in 5X SSC buffer was added to each sample and incubated at 600 rpm for 30 min at room temperature. Magnetic racks were used to separate the beads from unbound tRNA and the magnetic beads were washed once using 50 µL 1X SSC buffer and three times using 25 µL 0.1 X SSC buffer. Elution of the desired tRNA isoacceptor was carried out in 20 µL water for 3 min at 75°C. The samples were immediately placed into a magnetic rack, the supernatant was transferred into fresh tubes, and samples were directly prepared for nucleoside Liquid Chromatography-Tandem Mass Spectrometry (LC-MS/MS) analysis.

### Preparation for nucleoside LC-MS/MS

Total tRNA was diluted to a final concentration of 10 ng µL^-1^ and a final volume of 20 µL. Isolated tRNA isoacceptor samples were used as described in the respective methods section. RNA was digested to single nucleosides using a digestion master mix containing 2 U benzonase, 2 U alkaline phosphatase, and 0.2 U phosphodiesterase I in 5 mM Tris (pH 8) and 1 mM MgCl2-containing buffer. For nucleoside protection, 0.5 µg of pyrimidine deamination inhibitor tetrahydrouridine, 0.1 µg of purine deamination inhibitor pentostatin, and 1 µM of antioxidant butylated hydroxytoluene were added. The digestion mixture (total volume 35 µL) was incubated for 2 h at 37°C. 10 µL of LC-MS buffer was added after the digestion procedure.

For quantitative analysis, a calibration mixture was prepared using synthetic nucleosides. The calibration solutions ranged from 0.025 to 100 pmol for canonical nucleosides and from 0.00125 pmol to 5 pmol for modified nucleosides–pseudouridine ranged from 0.005 to 20 pmol. 10 µL of each sample and each calibration solution was injected into the LC-MS system for analysis. Additionally, 1 µL of nucleoside digested stable isotope labeled internal standard (SILIS) (50) was co-injected.

### LC-MS/MS of nucleosides

For quantitative mass spectrometry of nucleosides, an Agilent 1290 Infinity II equipped with a diode-array detector (DAD) combined with an Agilent Technologies G6470A Triple Quad system and electrospray ionization (ESI-MS, Agilent Jetstream) was used. The instrument was operated in positive-ion mode and dynamic multiple reaction monitoring (dMRM) for data acquisition. Further operating parameters: skimmer voltage of 15 V, cell accelerator voltage of 5 V, N2 gas temperature of 230°C and N2 gas flow of 6 L/min, N2 sheath gas temperature of 400°C with a flow of 12 L/min, capillary voltage of 2500 V, nozzle voltage of 0 V, nebulizer at 40 psi.

For separation prior to MS, a Synergi, 2.5 µm Fusion-RP, 100 Å, 100 x 2 mm column (Phenomenex, Torrance, California, USA) was used at a flow rate of 0.35 mL/min and a column oven temperature of 35 °C. Mobile phase A consisted of 5 mM aqueous NH4OAc buffer, brought to a pH of 5.3 with glacial acetic acid (65 µL/L). Mobile phase B consisted of organic solvent acetonitrile (Roth, Ultra-LC-MS grade). The gradient started at 100 % A for 1 min. B was then increased to 10% over a period of 4 min and subsequently increased to 40% over a period of 2 min. This condition was maintained for 1 min. Starting conditions (100% A) were restored over a period of 0.5 min and maintained for re-equilibration for 2.5 min.

### Data analysis of nucleoside LC-MS/MS

Raw data was analyzed using quantitative and qualitative MassHunter Software from Agilent. The signals for each nucleoside from dMRM acquisition were integrated along with the respective SILIS. The signal areas of nucleoside and respective SILIS were set into relation to calculate the nucleoside isotope factor (NIF, also see (51)), eq. 1:

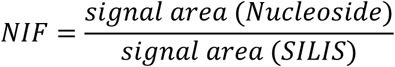

The nucleoside isotope factor was then plotted against the molar amount of each calibration and regression curves generated through the data points. The slopes represent the respective relative response factors for the nucleosides (rRFN) and enable absolute quantification of nucleosides. Calibration curves were plotted automatically by quantitative MassHunter software from Agilent. Molar amounts of samples nucleosides were calculated using the signal areas of the target compounds and SILIS in the samples and the respective rRFN, determined by calibration measurements. This step was performed automatically by quantitative MassHunter software. The detailed calculation is depicted in eq. 2:

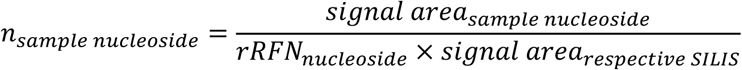

The molar amount of modified nucleosides was normalized to the molar amount of 1000 canonical nucleosides (eq. 3):

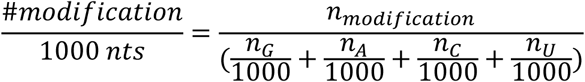

Statistical analysis was done using Excel and GraphPad Prism and statistical significance calculated by Welch’s t-test (p < 0.05 was considered significant).

### Quantitative reverse transcription PCR (RT-qPCR)

Prior to RT-qPCR analysis, complementary DNA (cDNA) was synthesized from 200 ng RNA templates using the Maxima H Minus Reverse Transcriptase kit (Thermo Fisher) following the manufacturer instructions. The cDNA concentration was quantified using the Qubit Flex Fluorometer (Thermo Fisher Scientific). RT-qPCR reactions were conducted using the Luna Universal qPCR Master Mix (New England BioLabs) according to manufacturer’s instructions. Each reaction contained 1 ng cDNA template. Technical triplicates for four to six biological replicates were performed for each experiment. The RT-qPCR program ran on QuantStudio 3 Real Time PCR System (Thermo Fisher), with the temperature profile in accordance with the Luna Universal qPCR Master Mix instructions. The threshold value (Ct) was determined by default settings. Relative expression was calculated using the 2^-ΔΔCT^ method (52) and was normalized to the expression values of *cox5* housekeeping gene (40).

### mRNA sequencing and analysis

For mRNA sequencing, CEA17 Δ*akuB^KU80^* and Δ*mod5* fungal samples (5x10^7^ spores) were cultured either for 24 h or for 16 h in 50 mL of AMM followed by an 8-h treatment with 80 ng/µL of 5-FC. RNA isolation and purification were carried out as explained above. The extracted RNA samples were subsequently submitted to Novogene (England) for quality control, library preparation, and mRNA sequencing. Sequencing was performed on the Illumina NovaSeq 6000 Sequencing System, using 150-bp paired-end sequencing. The raw data output, provided in FASTQ format by Novogene, was checked for quality, aligned to the A1163 fungal genome (Ensembl), and assessed for differential gene expression in accordance to (40). Briefly, pre-processing of raw reads, quality control and gene abundance estimation used GEO2RNaseq pipeline (v0.100.3, (53)) in R (version 3.6.3). Quality analysis was performed with FastQC (v0.11.5) before and after trimming. Read-quality trimming was done with Trimmomatic (v0.36) and reads were rRNA-filtered using SortMeRNA (v2.1) with a single rRNA database combining all rRNA databases shipped with SortMeRNA. Reference annotation was created by extracting and combining exon features from annotation files. Reads were mapped against reference genome of *A. fumigatus* (A1163, assembly GCA_000150145.1) using HiSat2 (v2.1.0, paired-end mode). Gene abundance estimation was done with featureCounts (v2.0.1) in paired-end mode with default parameters. MultiQC version 1.7 was used to summarize and assess quality of output from FastQC, Trimmomatic, HiSat, featureCounts and SAMtools. The count matrix with gene abundance data without and with median-of-ratios normalization (MRN, (54)) were extracted. Differential gene expression was analyzed with GEO2RNaseq. Pairwise tests were performed using four statistical tools (DESeq v1.38.0, DESeq2 v1.26.0, limma voom v3.42.2 and edgeR v3.28.1) to report *p*-values and multiple testing corrected *p*-values with false-discovery rate method q = FDR(p) used for each tool. Mean MRN values were computed per test per group including corresponding log2 of fold-changes. Gene expression differences were considered significant only if reported significant by all four tools (q ≤ 0.05) and |log2(fold-change [MRN based])| ≥ 1.

### Nano-tRNAseq

To perform Nano-tRNAseq, CEA17 Δ*akuB^KU80^* and Δ*mod5* fungal samples (5x10^7^ spores) were cultured for 24 h in 50 mL of AMM. The harvested mycelia were crushed with mortar and pestle under liquid nitrogen and subjected to RNA isolation as explained above. At least 20 µg per extracted RNA samples were subsequently submitted to IMMAGINA Biotechnology S.r.l. (Italy) for quality control, library preparation, and nanopore sequencing. The company performed sequencing and bioinformatic assessment according to (55) with slight modifications to facilitate sample multiplexing and the use of the *A. fumigatus* tRNA repertoire.

### Northern Blotting

Northern blots were performed to visualize tRNA expression patterns. The RNA probe was labelled using a DIG oligonucleotide tailing kit (Roche, Germany), as per the manufacturer’s instructions. 5-10 µg of DNase-treated RNA samples were denatured for 3 min at 70°C and separated on a 15% (w/v) TBE-urea gel (Novex®) with a low range ssRNA ladder (New England Biolabs, USA). The gel was stained with 1x SYBR gold dye (Thermo Fisher Scientific GmbH, Dreieich) and imaged with AlphaImager^TM^ (Biozym Scientific GmbH) imaging system. Subsequently, the RNA was transferred to a Roti®-Nylon plus membrane (pore size of 0.45 µM) with sufficient volumes of transfer buffer (10x SSC) for 4 h or overnight. Following capillary transfer, RNA-to-membrane-crosslinking was performed by UV-irradiation (2 min, E=1200 µJ/cm^2^) and the membrane was subjected to pre-hybridization for 1 h at 37-55°C (depending on probe) adding 20 mL of hybridization buffer (1 g dextran sulfate, 13.8 mL sterile water, 5 mL 20x SSC buffer, 10% SDS (w/v), 1 mL of 10x Western blocking reagent (Roche GmbH; Mannheim, Germany)). Hybridization was performed overnight after adding 25 pmol of the DIG-11-dUTP labelled probe. The membrane was washed at RT employing wash buffer I (1x SSC, 0.1% SDS) and wash buffer II (0.5xSSC, 0.1% SDS) for 5 min each. Afterwards, it was rinsed with 50 mL maleic acid buffer (0.15 M NaCl, pH 7.5- adjusted with NaOH pellets) for 5 min prior incubation with blocking solution (30 min, 5 rpm; 18 mL maleic acid buffer, 2 mL 10x Western Blocking reagent). The incubation was continued for another 30 min after addition of 0.5 µL of Anti-Digoxigenin-AP, Fab fragments (Roche, Mannheim, Germany). Unbound antibodies were eliminated by washing with 50 mL maleic acid buffer + 150 µL Tween 20 (Sigma Aldrich) for 15 min. Subsequently, the membrane was equilibrated by several washes with detection buffer (0.1 M Tris, 0.1 M NaCl, pH 9 – adjusted with HCl) and then incubated with CDP-Star substrate (Roche, Germany) for 5 min in darkness. The chemiluminescent signal was detected using the Chemiluminescence Imaging - Fusion FX SPECTRA (Vilber Lourmat, Collégien France). Potential reuse of the blots was executed by stripping with 0.5% SDS (w/v) for 1 h at 60°C, and re-hybridizing with a control probe.

### Southern Blot

Validation of target gene deletion or gene insertion in *A. fumigatus* were conducted by analytic DNA digestions and subsequent Southern Blot according to (40,56).

### Protein Extraction

Protein extraction for LC-MS/MS proteome analysis utilized the same hyphal samples as the mRNA sequencing. Equivalent amounts (∼ 50 mg) of mycelium were disrupted with mortar and pestle in liquid nitrogen, transferred into 1.5 mL screw-cap tubes and mixed thoroughly with 300 µL of freshly prepared lysis buffer (1% SDS (w/v), 150 mM NaCl, 100 mM TEAB (triethyl ammonium bicarbonate), 1 tablet Complete Ultra Protease Inhibitor Cocktail/PhosSTOP (Roche), ultrapure water). After addition of Benzonase nuclease (100 U) the samples were subjected to 30-min sonication (Bandelin Sonorex Super RK 510 H) at 37°C. To remove insolubilized debris, the samples were centrifuged (4°C, 15 min, 20000 x g) and the supernatant was further processed. Protein concentration was assessed using Merck Millipore Direct Detect System. 100 µg of protein was mixed with 100 mM TEAB, 2 µL reduction buffer (500 mM TCEP (tris(2-carboxyethyl) phosphine), 100 mM TEAB), and 2 µL freshly prepared alkylation buffer (625 mM 2-chloroacetamide dissolved in 100 mM TEAB). Afterwards, the samples were incubated at 70°C and 500 rpm in the dark for 30 min. Proteins were precipitated by MeOH/chloroform/water according to the protocol of Wessel and Flügge (57). Subsequently the pellet was dried in Concentrator plus (Eppendorf, Hamburg, Germany) and the protein precipitate was resolubilized in 100 µL of 100 mM TEAB in 5% (v/v) 2,2,2-trifluoroethanol and sonicated for 15 min. Next, Trypsin/LysC protease mix (2 µg/ µL, Promega) was added (1:25 protease:protein ratio) and incubated for 20 h at 37°C. Samples were afterwards dried in Concentrator plus (Eppendorf, Hamburg, Germany) and resolubilized in 30 µL 0.05% TFA, 2% acetonitrile in water. Ultrasonic bath treatment (15 min) and subsequent vortexing homogenized the samples. Filtration was then performed through 10 kDa MWCO spin filter (VWR), at 14 000 × *g* for 15 min at 8°C and the samples were transferred into high-performance liquid chromatography vials for further analysis.

### LC-MS/MS-based proteomics

LC-MS/MS analysis was performed according to (40), with slight adjustments mentioned in text below. Each biological sample was measured in three technical replicates. Analysis was conducted with Ultimate 3000 nano RSLC system coupled to Orbitrap Exploris 480 mass spectrometer (both Thermo Fisher Scientific, Waltham, MA, USA) with FAIMS. Peptide trapping took 5 min on an Acclaim Pep Map 100 column (2 cm x 75 µm, 3 µm) at 5 µL/min. This was followed by separation on an analytical Acclaim Pep Map RSLC nano column (50 cm x 75 µm, 2µm). Mobile phase gradient elution employed eluent A (0.1% (v/v) formic acid in water) mixed with eluent B (0.1% (v/v) formic acid in 90/10 acetonitrile/water). The following gradient was utilized: 0 min at 4 % B, 20 min at 6 % B, 45 min at 10 % B, 75 min at 16 % B, 105 min at 25 % B, 135 min at 45 %, 150 min at 65 %, 160-165 min at 96% B, 165.1-180 min at 4% B. Positively charged ions were generated at spray voltage of 2.2 kV with stainless steel emitter attached to Nanospray Flex Ion Source (Thermo Fisher). Quadrupole/orbitrap instrument operated in Full MS / data-dependent MS2 mode. Precursor ions were monitored at m/z 300-1200 at a resolution of 120,000 FWHM (full width at half maximum) using a maximum injection time (ITmax) of 50 ms and 300% normalized AGC (automatic gain control) target. Precursor ions with a charge state of z=2-5 were filtered at an isolation width of m/z 4.0 amu for further fragmentation at 28% HCD collision energy. MS2 ions were scanned at 15,000 FWHM (ITmax=40 ms, AGC= 200%) with three different compensation voltages being applied (-48 V, -63 V, -78 V).

### Protein database search

Tandem mass spectra were searched against the UniProt pan proteome database (2023/07/14) of *Aspergillus fumigatus* (https://ftp.uniprot.org/pub/databases/uniprot/current_release/knowledgebase/pan_proteome s/UP000002530.fasta.gz) using Proteome Discoverer (PD) 3.0 (Thermo) and the database search algorithms (threshold search engine scores in parenthesis) Chimerys (>2), Mascot 2.8 (>30), Comet (>3), MS Amanda 2.0 (>300), Sequest HT (>3) with and without INFERYS Rescoring. Two missed cleavages were allowed for the tryptic digestion. The precursor mass tolerance was set to 10 ppm and the fragment mass tolerance was set to 0.02 Da. Modifications were defined as dynamic Met oxidation, phosphorylation of Ser, Thr, and Tyr, protein N-term acetylation with and without Met-loss as well as static Cys carbamidomethylation. A strict false discovery rate (FDR) < 1% (peptide and protein level) was required for positive protein hits. The Percolator node of PD3.0 and a reverse decoy database was used for q-value validation of spectral matches. Only rank 1 proteins and peptides of the top scored proteins were counted. Label-free protein quantification was based on the Minora algorithm of PD3.0 using the precursor abundance based on intensity and a signal-to-noise ratio >5. Normalization was performed by using the total peptide amount method. Imputation of missing quan values was applied by using abundance values of 75% of the lowest abundance identified per sample. Differential protein abundance was defined as a fold change of >2, pvalue/ABS(log4ratio) <0.05 (pvalue/log4ratio) and at least identified in 2 of 3 replicates of the sample group with the highest abundance.

### Codon Usage Analysis

Codon counts per coding sequence were determined from the *A. fumigatus* A1163 ASM15014v1 cds fasta file available on Ensembl Fungi using R version 4.3.0 (2023-04-21) and the Biostrings Package (58). Plots and correlation analyses were produced as described in the text using R package ggplot2 version 3.4.4 (59).

### tRNA gene distance calculation

The distance between mRNA and nearest tRNA gene was determined by extracting coordinates from the *A. fumigatus* A1163 ASM15014v1.57 gff3 file (Ensembl Fungi) and comparing annotated tRNA and mRNA features using bedtools (v2.31.0) *closest* (60). mRNA elements on short contigs lacking tRNAs were removed from the downstream analyses.

### Data and statistical analysis

Statistical analyses were performed in GraphPad Prism 10 or using R version 4.3.0 (2023-04- 21) in R Studio Version 2023.06.0+421 (2023.06.0+421) as described in detail in the specific methods sections or figure legends as appropriate.

## RESULTS

### Deletion of *mod5* confers 5-fluoroorotic acid and 5-fluorocytosine resistance

*A. fumigatus* relies on rapid growth and a coordinated stress response to establish infections in the human host (61). As orthologs of Mod5 serve as regulatory hubs for stress response in bacteria, we sought to test for similar importance of *mod5* in a eukaryotic pathogen (21,22,62).

Using FungiDB, we identified *AFUB_093210* (hereafter called *mod5*) encoding a putative IPTase ortholog in *A. fumigatus* (63). Comparison of amino acid sequences of known Mod5- orthologs from *E. coli* (MiaA), *Homo sapiens* (TRIT1), *S. cerevisiae* (Mod5), and *S. pombe* (Tit1) revealed the conservation of all relevant domains (**Fig. S1**) (64,65). We generated a Δ*mod5* strain and subjected it together with the wild-type to tRNA or total RNA isolation and subsequent LC-MS/MS and Nano-tRNAseq analysis to verify the lack of the corresponding i^6^A modification and identify potential Mod5 targets, respectively (**Fig. 1A,B; Fig. S2,3; Table S3- 5**). We observed a complete absence of the modification in total tRNA and confirmed tRNA^Tyr^GΨA, tRNA^Cys^GCA, tRNA^Ser^, and tRNA^Ser^ as Mod5 modification targets in *A. fumigatus* (**Fig. 1A,B; Fig. S3**), with mttRNA^Tyr^GΨA, mttRNA^Cys^GCA, and mttRNA^Sup^ identified as i^6^A-modified among the mitochondrial tRNA pool (**Fig. 1B; Fig. S3**). Following knockout validation, droplet assays were conducted to assess *Δmod5* growth under several stress situations (**Fig. 1**). Compared to the wild type, the *mod5* knockout displayed a slightly reduced colony size on AMM plates under standard growth conditions, which was not visible on rich malt agar plates (**Fig. 1A**).

**Figure 1.**
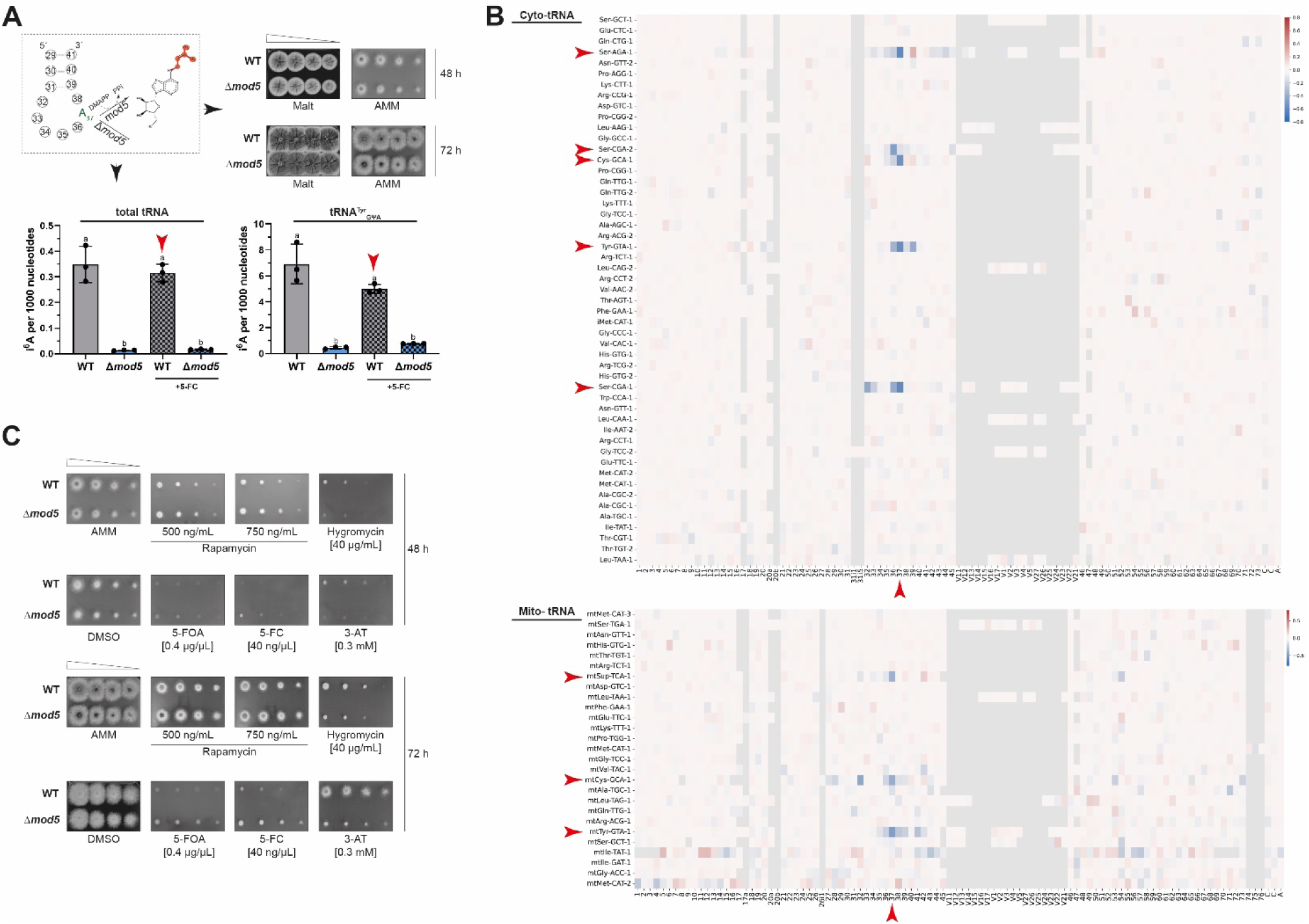
Deletion of *mod5* (*AFUB_093210*) in *A. fumigatus* results in increased resistance against 5-FC and 5-FOA. (A) Model displaying the i^6^A37-modification process depending on Mod5. The right panel shows droplet growth assays for the wild-type and Δ*mod5* strains grown on malt and AMM agar for 48 h and 72 h. The lower panel displays the LC-MS/MS based quantification of i^6^A37 levels in total tRNAs/tRNA^Tyr^GΨA isolated from the wild type and Δ*mod5* under non-stressed and 5-FC-stressed conditions. Statistical significance was calculated with Welch’s t-test and indicated by small letters. A *P*-value ≤ 0.05 was considered significant. (B) Summary heatmaps of the Nano-tRNAseq experiment displaying basecalling errors of Δ*mod5* relative to the wild type for the cytoplasmic and mitochondrial tRNA pool. The x-axis provides the nucleotide position while the y-axis indicates all tRNA isoacceptors listed according to their abundance (top: most abundant to bottom: least abundant). Red arrows indicate position 37 and all tRNA species displaying a base calling error at this site. (C) Strains were spotted and cultivated on AMM agar plates containing either 500 ng/mL or 750 ng/mL rapamycin, 40 µg/mL hygromycin, 0.4 µg/µL 5-FOA, 40 ng/µL 5-FC, or 0.3 mM 3-AT. Wild type and Δ*mod5* grown on AMM (without DMSO) agar plates served as controls. Documentation of the different assays was carried out after 48 h and 72 h. Data are representative images from three biological replicates.

Treatment of Δ*mod5* with 3-amino-triazole (3-AT) resulted in severe growth defects visible after 48 h of incubation and even more prominent after 72 h (**Fig. 1C**). 3-AT is known to inhibit histidine biosynthesis, which decreases aminoacylation of tRNA^His^ and activates the amino acid starvation responses known as cross-pathway control (CPC) and/or general amino-acid control (GAAC; (66–69)). Since different anticodon stem-loop tRNA modification defects result in premature CPC/GAAC induction (70), application of 3-AT may also trigger this response in Δ*mod5*. Hygromycin treatment also affected the growth of the Δ*mod5* knockout, likely indicating decreased frame maintenance during translation. Inhibition of TOR using two different concentrations of rapamycin (500 ng/µL and 750 ng/µL; (38)) resulted in a growth reduction of both the wild type and Δ*mod5* compared to the AMM control. Growth on agar plates containing either 5-FOA or 5-FC, both known to disrupt DNA and RNA stability (71), was severely impeded for the wild type, whereas the Δ*mod5* strain surprisingly appeared resistant to those antifungal drugs. This resistance was even more pronounced after 72 h, in contrast to other fungal systems where IPTase knockouts displayed increased susceptibility under similar stress situations (72,73). The MIC for Δ*mod5* was established with minor modifications to the EUCAST guidelines ((74); **Table S6**) as 0.4 -0.5 µg/µL for the wild type and roughly 5x fold higher for Δ*mod5* at 2-3 µg/µL.

Interference with β-(1,3)-d-glucan biosynthesis using caspofungin (0.5 µg/mL and 8 µg/mL) resulted in strongly reduced growth of the *mod5* knockout, whereas blocking ergosterol biosynthesis (75) with tebuconazole, itraconazole, and voriconazole using three independently generated clones of the deletion strain displayed variable decreases in growth compared to the wild type (**Fig. S4**). Although rewiring of the i^6^A37 substrate dimethylallyl pyrophosphate (DMAPP) towards ergosterol biosynthesis and subsequent azole resistance was reported upon abolishment of the tRNA modification in *S. cerevisiae* (37), we could not confirm this for *A. fumigatus*. This implied that the absence of Mod5 resulted not in a general resistance against antifungal drugs but instead affects the susceptibility against the fluorinated pyrimidine analogues, 5-FC/5-FOA. In conclusion, both droplet and MIC assays confirm resistance of the *mod5* knockout against the antifungal agents 5-FC and 5-FOA.

### Antifungal resistance of Δ*mod5* is conferred independently of the canonical *fcyB* axis

We next assessed the transcriptome and proteome of the knockout strain to better understand the 5-FC resistance phenotype of Δ*mod5* and explore the contribution of i^6^A37 loss to regulation of gene expression. Previous studies found changes in the transcriptome and proteome attributable to tRNA modification defects that resulted in the induction of starvation and stress response pathways (16,38,70,76). We hypothesized that similar changes give Δ*mod5* an advantage in coping with 5-FC. For both experiments we used mycelium of the wild type and the knockout from liquid cultures that were grown either for 24 h non-stressed or for 16 h of normal growth followed by 8 h with 5-FC. The material obtained was split and subjected to either RNA or protein isolation for poly-A enriched RNA-seq or label-free quantification (LFQ) proteomics, respectively. Notably, since 5-FC is converted into 5-fluorouracil (5-FU) and then 5-fluorouridine triphosphate (5FUTP) before incorporation into de-novo synthesized transcripts (77), we also observed an expected minor decrease in RNA-stability for our 5-FC-treated samples. The sequencing of each strain under both conditions was used to assess expression changes under non-stressed conditions (Δ*mod5* vs WT), 5-FC-stressed and non-stressed conditions (Δ*mod5*+5-FC vs Δ*mod5*, WT+5-FC vs WT), and 5-FC treatment only (Δ*mod5*+5- FC vs WT+5-FC) (**Fig. 2A,B**; **Table S7,8, Fig. S5**). From the RNA-seq, we recognized that the *mod5* knockout displayed notable changes to the transcriptome under standard growth conditions, inducing 156 genes and repressing 44 (**Fig. 2A**; **Table S8**).

**Figure 2.**
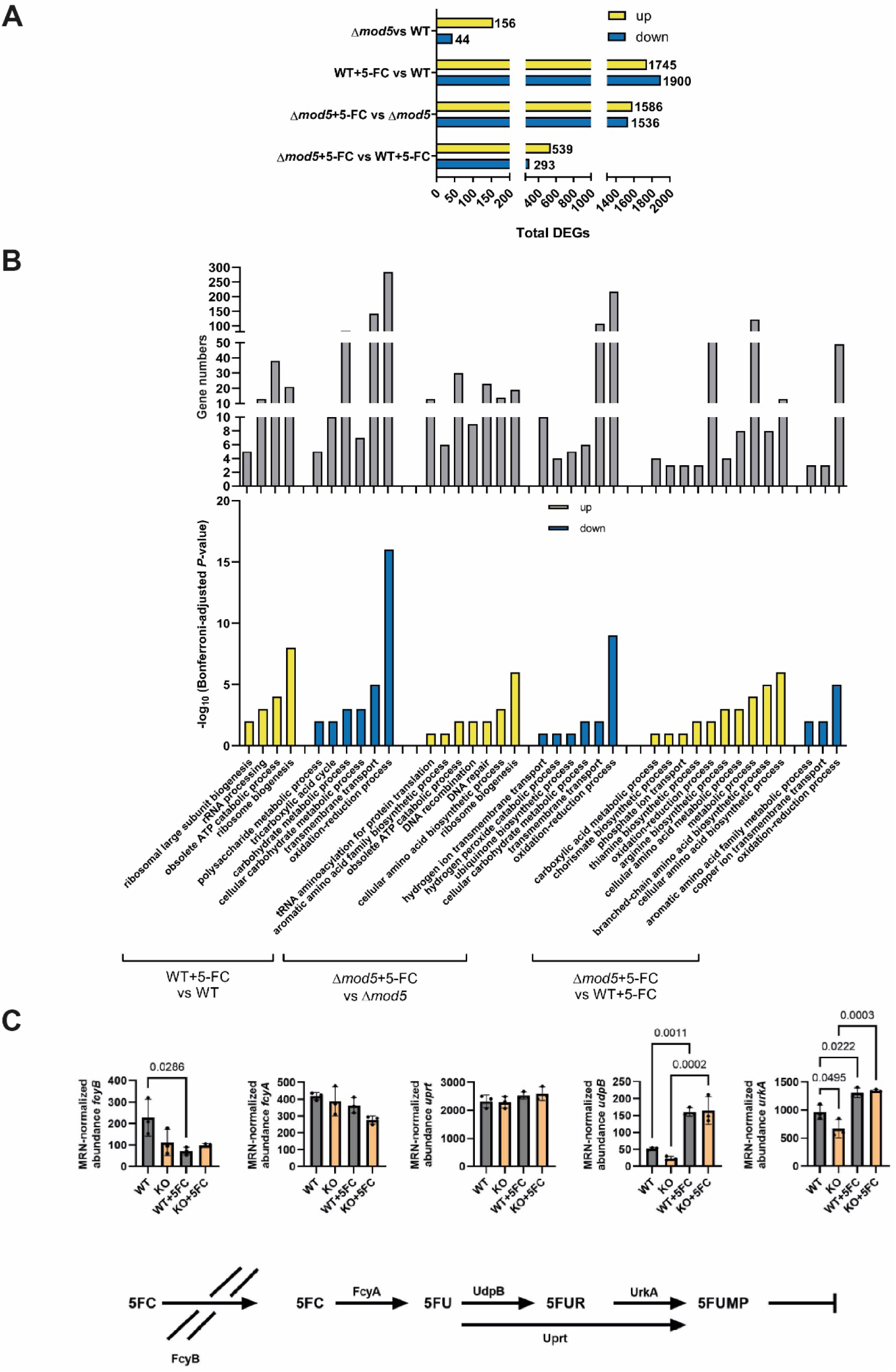
Wild type and Δ*mod5* transcriptome differences are inflated upon 5-FC treatment. (A) RNA-seq-identified significant genes (*P*≤0.01) with a log2 fold change of at least 1 (log2 FC≥1) are given for the indicated comparisons between strains (Δ*mod5*/wild type) and growth conditions (-/+ 5-FC). Induced genes are given in yellow and suppressed ones in blue. (B) GO- terms for up- (yellow) and down-regulated (blue) genes among the different comparisons are listed according to their Bonferroni-adjusted *P*-values (-log10) together with the respective gene numbers (grey). (C) The model displays the canonical 5-FC salvaging pathway in *A. fumigatus* with all participating proteins. Normalized MRN-abundances of *fcyB*, *fcyA*, *uprT*, *udpB*, and *urkA* are given for the wild type and Δ*mod5* under non-stressed and 5-FC stressed conditions. Significance was calculated with one-way ANOVA and Šídák’s multiple comparisons test.

Gene ontology (GO) analysis of the differentially expressed genes (DEGs) yielded an enrichment in the “cellular compartment” category for genes encoding membrane-localized transport proteins (26 induced and 11 suppressed), hinting at changes in membrane-trafficking (**Table S9**) (78). Further inspection of the altered genes revealed expression changes of different aspects of metabolically relevant genes that included secondary and carbohydrate metabolism (**Table S10,11**). These transcriptional alterations together might suggest a general mis-sensing of the nutrient level in the environment and an adaptation process of Δ*mod5* in line with reports of multiple other tRNA modification mutants (16,38,70,76). Next, we asked if any genes of the canonical 5-FC salvaging pathway were also differentially regulated in unstressed conditions. The uptake of 5-FC is mediated by the cytosine permease FcyB and is converted stepwise to 5-FU by the cytosine deaminase FcyA and uracil phosphoribosyltransferase Uprt, before feeding into RNA-biosynthesis to affect transcript stability, disrupt translation, and inhibit production of thymidylate synthase to impede DNA-synthesis (**Fig. 2C; Table S7,8**) (77,79,80). In addition to the three canonical proteins it was recently established that two other proteins, UrkA and UdpB, are also involved in *A. fumigatus* 5-FC metabolism (80). We observed a significant, but minor reduction in the expression of *urkA* and no change in *udpB* in Δ*mod5* compared to wild type under non-stressed conditions.

To assess the response to 5-FC stress more broadly, we interrogated the stress-dependent changes observed by RNA-seq more closely. As expected, treatment of the wild type and *mod5* knockout with 5-FC resulted in dramatic changes of both transcriptomes. The wild type displayed 3,645 DEGs (1,745 induced genes, 1,900 suppressed genes), while Δ*mod5* showed expression changes of 3,122 genes (1,586 induced genes, 1,536 suppressed genes) (**Fig. 2A**; **Table S8**) after 5-FC stress. We again checked for significant changes in 5-FC metabolism genes but only observed a slight upregulation of *udpB* and *urkA* in the *mod5* knockout and the wild type (**Fig. 2C, Table S7,8**). It is then unlikely, that the canonical 5-FC salvaging pathway is involved in the observed Δ*mod5* resistance phenotype. Consequently, we analysed the substantial changes observed for both strains to understand the pathways that might contribute to the resistance phenotype. A GO-term analysis (biological processes) revealed that both the wild type and the knockout displayed similar transcriptional changes embodied by an induction of translation-relevant genes (**Fig. 2B**; **Table S12,13**). Similar effects upon 5-FC treatment have been observed in other pathogenic fungi like *C. albicans* and *Candida glabrata*, probably indicating a compensatory effect to cope with increased translational defects (81–83). Both strains appeared to activate DNA damage response pathways to cope with corresponding 5- FUTP-related damage, but the effect only reached statistical significance in the knockout, likely due to higher variability in the wild-type results (**Fig. 2B**; **Table S12,13**)(77). Moreover, various GO-terms implied differential regulation of transmembrane transport proteins (e.g., “transmembrane transport”) for both strains, in line with changes of transmembrane transporter regulation as well as DNA repair/recombination activity reported in other 5-FC treated fungal species like *S. cerevisiae* (81,84,85). Remarkably, additional shifts were executed by both strains in carbohydrate metabolism and redox regulation (**Fig. 2B**; **Table S12,13**), including suppression of genes relevant for energy generation, production of energy reserves like glycogen or trehalose, and membrane-associated electron transport and energy conservation. All these transcriptional changes point to a potential alteration of mitochondrial energy generation.

Finally, we performed a transcriptome comparison of both 5-FC-challenged wild-type and Δ*mod5* strains to further explain the observed resistance phenotype. GO-term analysis intriguingly revealed that the tRNA modifier knockout strongly induces metabolically relevant genes that are involved in amino acid biosynthesis (**Fig. 2B**; **Table S14**). Similar to other fungal species, the amino acid starvation response in *A. fumigatus* is controlled by the transcription factor CpcA and the cross-pathway control (CPC) system (86). This virulence-relevant pathway is canonically triggered by a decline of amino-acylated tRNAs, which in the case of 5- FC treatment is known to increase rapidly due to the incorporation of 5-FUTP (77). Compensation for this effect is also evident in the RNA-seq dataset (e.g. Δ*mod5* +/- 5-FC, “tRNA aminoacylation for protein translation”; **Fig. 2B; Table S13**) resulting in the induction of 13 and 10 aminoacyl tRNA-synthetases (aaRS) in the knockout and wild type, respectively. Nevertheless, the induction of CPC upon 5-FC treatment appears to be stronger in Δ*mod5* than the wild type, likely due to premature activation as previously reported (16,38,70,76).

### Tyrosine serves as the most dominant Mod5 target codon

The link between tRNA modification and translation led us to hypothesize that the proteome would display differences compared to the transcriptome in the knockout. We performed LC-MS/MS based label-free quantification on the wild-type and Δ*mod5* strains using proteins collected simultaneously from the same exact samples as in the RNA-seq experiment. We identified 4,908 proteins common in all conditions, including both strains under non-stressed and 5-FC stressed conditions (WT: 5,551 proteins, Δ*mod5*(KO): 5,556 proteins, WT+5-FC: 5,117, Δ*mod5*(KO)+5-FC: 5,337 proteins; **Table S15**). Due to the stringency of the Benjamini-Hochberg analysis in correcting for multiple comparisons and the potential of missing many other true differentially abundant proteins, we performed an analysis using the two-tailed Welch’s *t*-test (87). Using this approach, we identified many proteins differing significantly in abundance by at least two-fold (**Fig. 3A**; **Fig. S6A-D**). Consistent with the transcriptome dataset, we saw comparable effects for the induced/suppressed proteins in the Δ*mod5* and wild-type strains upon non-stressed and 5-FC stressed conditions (Δ*mod5* vs WT: 69 induced, 142 suppressed; WT+5-FC vs WT: 179 induced, 387 suppressed; Δ*mod5*+5-FC vs Δ*mod5*: 201 induced, 187 suppressed; Δ*mod5*+5-FC vs WT+5-FC: 12 induced, 14 suppressed; **Fig. 3A**).

**Figure 3.**
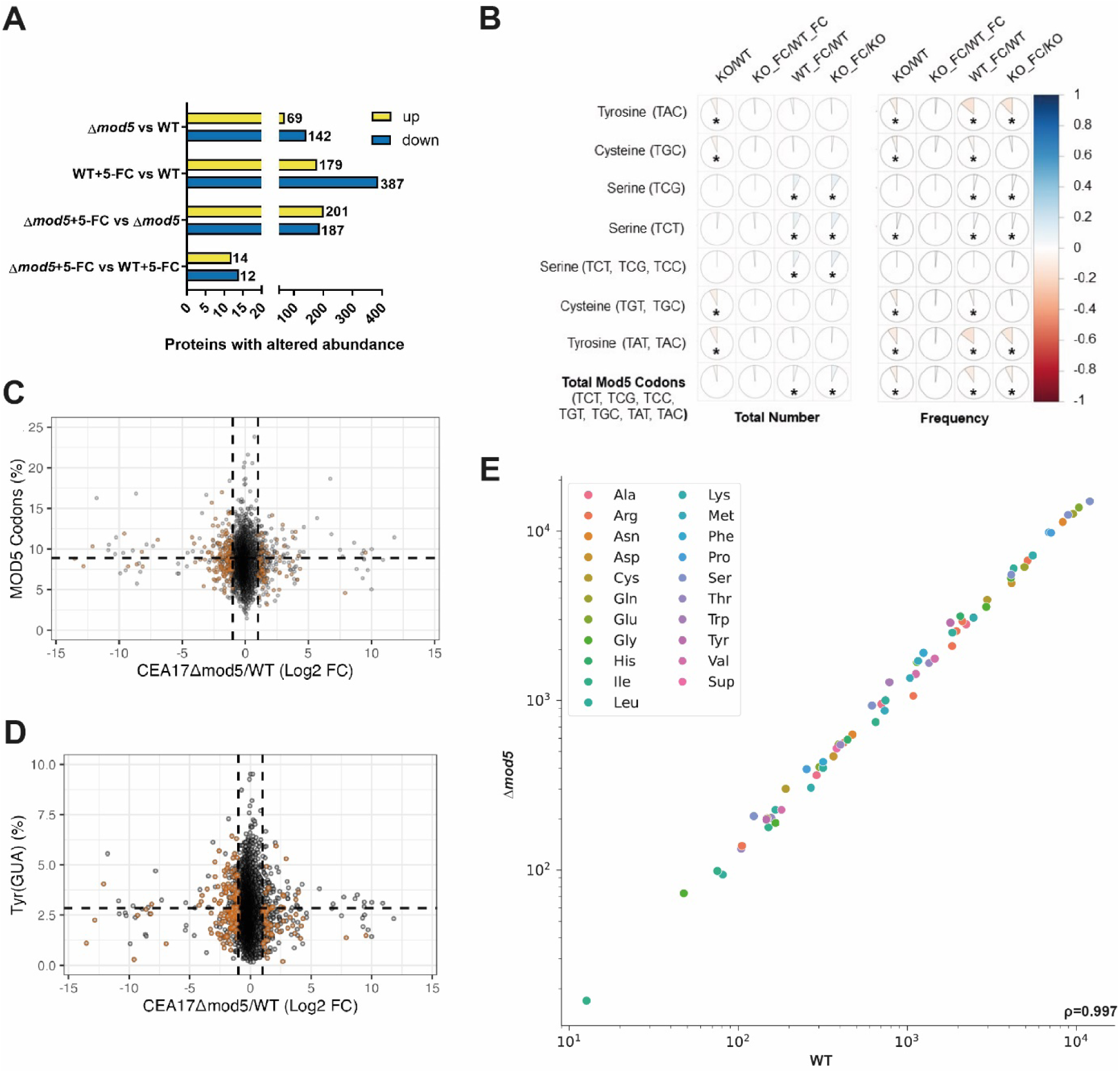
Proteome changes mirror transcriptional alterations elucidated for the wild type and Δ*mod5* under non-stressed and 5-FC stressed conditions. (A) LC-MS/MS-identified proteins with a log2FC≥2 and *P*≤0.05 are given for the indicated comparisons between strains (Δ*mod5*/wild type) and growth conditions (-/+ 5-FC). Proteins with increased abundance are given in yellow and decreased in blue. (B) Spearman’s rank correlation coefficients were determined for the number and frequency of Mod5-modified codons and select codon combinations (see **Table S16**) in coding sequences compared to the Log2FC values for the indicated comparisons determined in the proteomics analysis (**Table S17-20**). Blue and red imply a positive or negative correlation. (C,D) Plots display relative protein abundances (cut- off given in A) of the different comparisons conducted for Δ*mod5* and wild type upon different conditions (+/- 5-FC) versus Mod5-dependent codon-usage per open-reading frame (**Table S16-20**). Orange dots indicate samples with *P*≤0.05 and Log2FC >|1|in the corresponding proteomics comparison. To be included in (B-D), a protein had to be identified in at least 2 biological replicates of one of the tested conditions. (E) The tRNA abundances are displayed in a scatter plot correlating the Nano-tRNAseq data from all replicates of Δ*mod5* and wild type. The Spearmańs correlation coefficient (ρ) indicates the correlation strength.

To determine whether Mod5 influenced the general relationship between transcript expression and protein abundance, we plotted Z-scored protein abundances against Z-scored transcriptome data for the wild-type and Δ*mod5* strains (**Fig. S6E,F**). As expected, we observed a clear correlation between the proteome and transcriptome in the wild type, but also somewhat surprisingly for the knockout, suggesting that *mod5* deletion did not overtly skew translation. Both analyses resulted in moderate positive correlations as demonstrated by Pearson-correlation coefficients of (r) = 0.59 for the wild type and (r) = 0.58 for the Δ*mod5*. The plots are overlaid with the frequency of Mod5 target codons, with orange representing coding sequences more than one standard deviation above the average Mod5-target codon frequency of 0.089 (±0.024). There were no obvious differences between the different groups, suggesting that the linear relationship between the transcriptome and proteome was not grossly disrupted by *mod5* deletion.

We next assessed individual genes more specifically by assessing potential Mod5 modification-tuneable transcripts (MoTTs) (88,89). MoTTs are transcripts whose translation is regulated by tRNA modifications. We rationalized that the deletion of *mod5* and ablation of i^6^A37 might manifest in the proteome as Mod5-dependent MoTTs. Since we identified the Mod5 tRNA-targets for *A. fumigatus* by Nano-tRNAseq (**Fig. 1B; Fig. S3**), we focused on tRNA^Tyr^GΨA, tRNA^Cys^GCA, tRNA^Ser^, and tRNA^Ser^, which match highly conserved codons from the literature (16). Pearson correlation coefficients were calculated for each codon number (number of codons per coding sequence (CDS)) and respective codon frequency (specific codon / total codons in a CDS) compared to protein abundance (**Fig. 3B; Fig. S7,8; Table S16**). A negative correlation suggested a tRNA modification was required for proper translation, whereas a positive correlation suggested Mod5 activity decreased translation. We included comparisons of 5-FC treated and untreated samples to aid in finding codon-dependent deregulation (i.e. MoTTs) of specific proteins (**Fig. 3B-D; Fig. S7,8**). Most predicted Mod5 codons exhibited small, but significant negative correlations, with serine codons being the notable exception (16), likely due to the very high relative levels of serine tRNA in the fungus (**Fig. 3E**). A few unexpected codons did show slight positive (e.g., arginine) or negative correlations (e.g., tryptophan, histidine, proline), which may warrant further investigation in the future. The tyrosine codons (and to a lesser extent cysteine and tryptophan codons) were negatively correlated with protein abundance in the knockout under normal growth conditions, but no major differences were observed during 5-FC stress, suggesting that additional mechanisms may be involved in promoting antifungal resistance to 5-FC beyond tRNA modification function alone. We conclude that the changes observed for the proteome closely resemble those detected for the transcriptome and are apparently mildly attributed to the identified Mod5-dependent MoTTs as seen for the correlation with the modification of tyrosine- and cysteine-recognizing tRNAs.

### 5-FC treatment triggers cross-pathway control in *A. fumigatus*

The transcriptome and proteome datasets both indicated a substantial activation of genes involved in amino acid biosynthesis upon loss of *mod5* and/or 5-FC treatment (**Fig. 2B**, **Table S7,8**,**17-20**). Many of these signal cascades are known to be part of the evolutionarily conserved CPC system that is regulated by the transcription factor CpcA (orthologue of Gcn4; (86)). CpcA and Gcn4 are both controlled by various translational and transcriptional regulatory layers, whereas the *cpcA* gene bears an additional auto-regulatory mechanism amplifying its own expression (90). We made use of this feature by applying quantitative reverse-transcription PCR (RT-qPCR) to measure the expression of *cpcA* and one of its client genes *argB* under standard AMM-growth conditions of the wild-type and Δ*mod5* strains (**Fig. 4**, non-shaded bars) (90). In agreement with other reports from tRNA modification mutants, the deletion of *mod5* in *A. fumigatus* resulted in a similar premature activation of the CPC by inducing the expression of *cpcA* and *argB* 1.5-fold (70). To elucidate if Δ*mod5* also altered *cpcA*/*argB* induction upon CPC-activation, we quantified expression in 3-AT treated mycelium, a known canonical trigger of this pathway (90). The two strains induced *cpcA* and *argB* as expected, although the wild type displayed a 5.4-fold change and Δ*mod5* only a ≈4-fold change in comparison to the respective non-induced mRNA levels (**Fig. 4A**, shaded bars). When we repeated the experiment treating the wild type and knockout with 5-FC, we detected an induction of *cpcA* and *argB* in both strains (wild type: ≈8.5-fold and ≈2-fold, Δ*mod5*: ≈5-fold and ≈8-fold) in line with our transcriptional analysis (**Fig. 4B**, shaded bars). Surprisingly, while the induction of *cpcA* in the wild type was stronger than in Δ*mod5*, the *argB* level was substantially higher in the knockout. This suggests that either CpcA activity is differentially modulated in the Δ*mod5* strain, or more compellingly, that alternative transcription factors are activated to control *argB* expression in the absence of Mod5.

**Figure 4.**
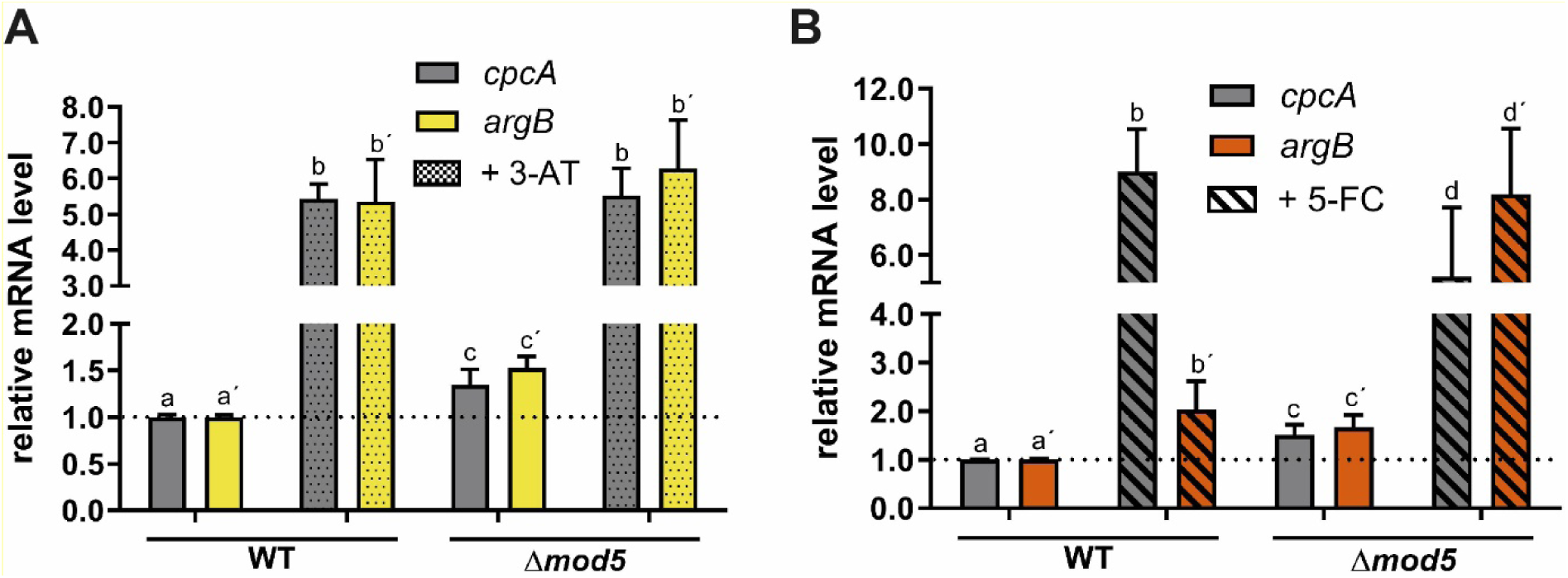
Treatment with 5-FC and 3-AT induce *cpcA* expression but results in differential induction of downstream client gene *argB*. Wild-type and Δ*mod5* mycelial balls were harvested after 24 h cultivation in AMM media and subjected to RNA isolation, cDNA synthesis and qPCR to quantify *cpcA* and *argB* mRNA level. Additionally, mycelial balls of the respective strains were treated either with 3-AT (A) or 5-FC (B) and quantified for the indicated genes. The quantifications of *cpcA*/*argB* were normalized to the *cox5* level (52). The calculated *P*-values are as follows: 0.05-0.01 (a-c; a’-c’; b-d); <0.01 (a-b; a’-b’; c-d; c’-d’; b’-d’). Statistical significance was calculated using a two-way ANOVA test on log10 transformed data. Pairwise comparisons were carried out using the Holm Sidak method. Small letters indicate statistical significance with the apostrophe (‘) sign differentiating between the genes (*cpcA*, *argB*). The *P*-value ≤ 0.05 was considered as significant.

### CpcA-driven induction of Major Facilitator Superfamily (MFS) transporter *nmeA* results in decreased antifungal activity in Δ*mod5*

We hypothesized that activation of the CPC pathway primes the tRNA modifier knockout to better cope with 5-FC treatment by activating appropriate resistance genes/mechanisms. In the literature, *A. nidulans nmeA (Ani_nmeA)*, a gene that encodes for a nucleobase/nucleoside exporter of the Major Facilitator Superfamily (MFS), is known to confer 5-FC resistance when overexpressed (91). Interrogating the RNA-seq data for any induced orthologues of Ani_*nmeA*, we found the ortholog *AFUB_005530* (renamed *nmeA*) to be slightly upregulated in the *mod5* knockout under standard growth conditions (**Fig. 5A,B; Table S8**). Treatment with 5-FC increased expression of *nmeA* in both the knockout and the wild type, although Δ*mod5* displayed significantly higher levels (**Fig. 5A,B; Table S8**). Additional *nmeA* orthologues were predicted in *A. fumigatus* according to FungiDB (OrthoMCL) that may also contribute to the export of 5-FC/byproducts upon CPC induction. Analysis of the RNA-seq data for the eight other genes only revealed a similar expression pattern for two other candidate genes *AFUB_043340* and *AFUB_028350* under the tested conditions (**Fig. 5C; Table S8**). Due to the potential CpcA-dependent activation, we assumed that *nmeA* and other orthologs might contain the *Aspergillus* genus-conserved CpcA binding site TGASTCA (S=G or C, (92)) within their promoters. All 9 gene promoter regions were investigated for CpcA binding sites and indeed, *nmeA*, *AFUB_043340*, and *AFUB_028350* were the only ones with such a sequence, in contrast to the *A. nidulans nmeA* promoter region (**Fig. 5**, black boxes), which had only one CpcA binding site in a unique location. Matching the RNA-seq reads of *nmeA* to its promoter region revealed that the potential CpcA recognition sequence resides close to the transcription start site (TSS; **Fig. 5B**, red arrows and boxes), suggesting that CPC-regulated expression of this exporter may result in a more proximal TSS and shorter untranslated region (UTR).

**Figure 5.**
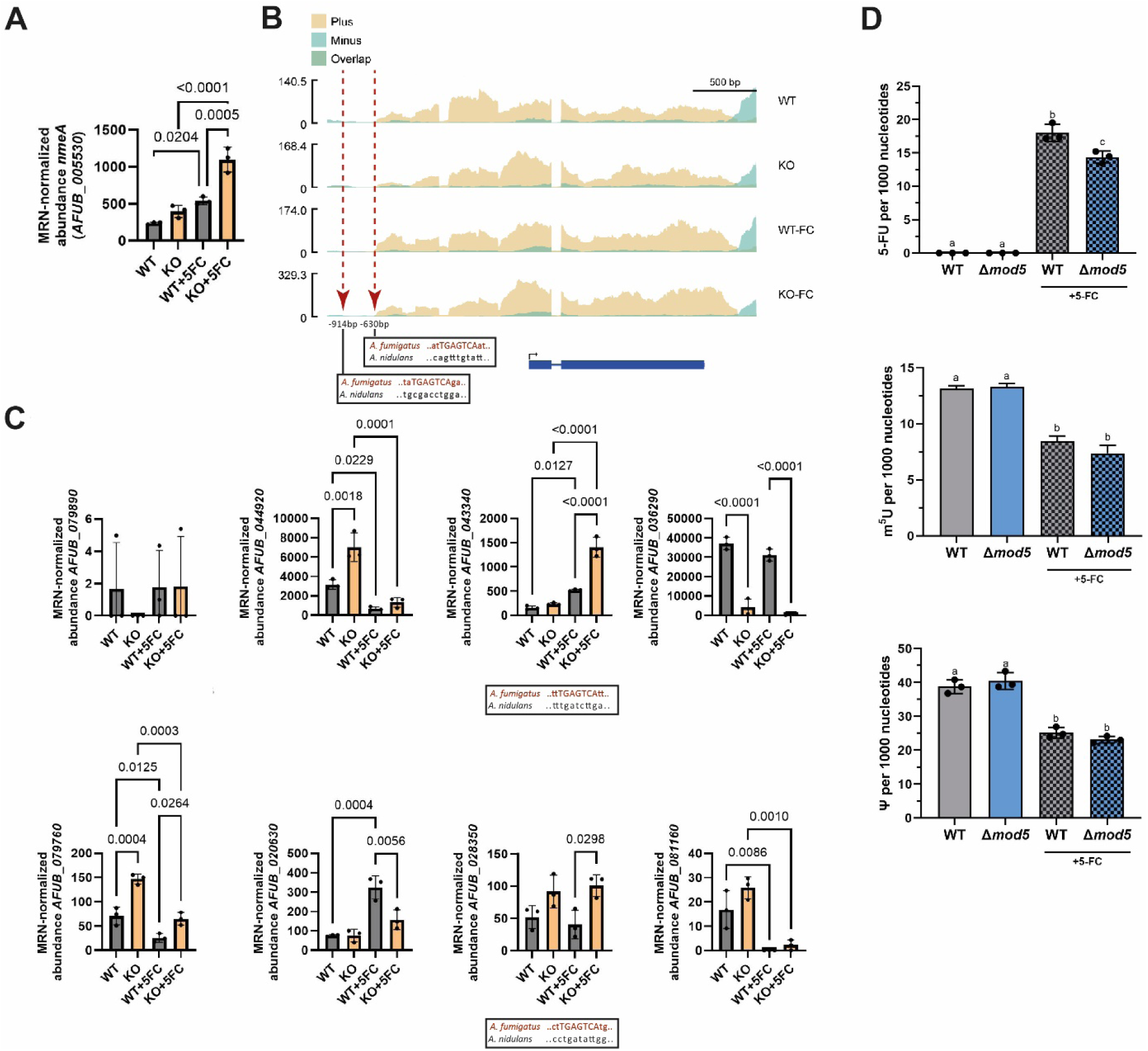
Ablation of i^6^A results in the CPC-driven induction of MFS exporter *nmeA* and decreases the 5-FU level in Δ*mod5*. (A) mRNA-seq expression data of *nmeA* (*AFUB_005530*) displayed for the wild type and Δ*mod5* upon non-stressed and 5-FC stressed conditions. (B) Obtained sequencing reads are plotted against the gene locus of *nmeA*. The red arrows and black boxes indicate found conserved CpcA binding sites. (C) Eight additional orthologues of *nmeA* are indicated with their expression values assessed by mRNA-seq in both strains and the given conditions. *AFUB_043340* and *AFUB_028350* contain putative CpcA binding sites in their promoter region (black boxes). (D) The LC-MS/MS based quantification of 5-FU, m^5^U and Ψ levels in total tRNAs isolated from the wild type and Δ*mod5* under non-stressed and 5- FC-stressed (checked bars) conditions are given. Statistical significance calculation was performed using Welch’s t-test and is indicated by small letters. A *P*-value ≤ 0.05 was considered significant. Statistics for (A) and (C) were conducted with one-way ANOVA and Šídák’s multiple comparisons test.

We hypothesized that premature and 5-FC-induced upregulation of *nmeA* (and potentially *AFUB_043340* and/or *AFUB_028350*) in Δ*mod5* would contribute to the resistance phenotype, as evidenced by reduced 5-FU incorporation into transcripts and interference with uridine- directed modifications in e.g., tRNAs, in the knockout compared to the wild type. To explore this idea, we quantified total transfer RNAs by LC-MS/MS for their 5-FU content and levels of i6A, Ψ, and 5-methyluridine (m^5^U), the latter two modifications known to be sensitive to fluorouracil treatment (93,94). The quantification of wild-type and knockout samples revealed a strong increase of 5-FU levels upon 5-FC treatment but a clear significant difference regarding transcript-incorporation, with Δ*mod5* displaying lower fluorouridine amounts (**Fig. 5D**, upper panel, **Table S5**). Correspondingly, when assessing Ψ and m^5^U levels, a 5- FC-dependent decrease of both modifications was visible in the wild type and knockout, but interestingly, no significant difference was observed between both strains (**Fig. 5D**, middle and bottom panel, **Table S5**). A similar event was visible upon measuring the i6A-levels in the wild-type total tRNA and tRNA^Tyr^GΨA, displaying an obvious but not significant decrease of the modification upon 5-FC treatment (**Fig. 1A, Table S5**), hinting that the fungus may naturally vary Mod5 levels or activity in response to stress. These results suggest that *A. fumigatus* adapts to the 5-FC threat in a CPC-dependent manner to compensate for potential damage by increasing expression of the *nmeA* exporter and reducing the available pool of 5-FC/5-FU incorporated into (t)RNA transcripts in Δ*mod5*.

### Deletion of *nmeA,* but not tRNA^Tyr^GΨA overexpression, reverses 5-FC resistance of Δ*mod5*

The resistance of Δ*mod5* to the antifungal agent 5-FC is likely CPC driven, as the deletion alone resulted in transcriptome and proteome alterations consistent with induction of CpcA- client genes under normal growth conditions (**Fig. 2-5**). Similar changes have been reported for other tRNA modification mutants, and in each case the phenotypes were rescuable by the overexpression of the respective, majorly affected tRNA (38,39,70,76). We hypothesized that resistance was attributed to the modification defect of individual or multiple target tRNAs (**Fig. 1, Fig. S3**) and corresponding translational impedance. We attempted to revert the 5-FC phenotype by providing an excess of the Mod5 tRNA target tRNA^Tyr^GΨA, which encodes the codon UAC with the highest frequency among the downregulated proteins in our proteomics (**Fig. 1, Fig. S3; Table S16**) (16) and is the only existing iso-decoder for tyrosine in *A. fumigatus* according to GtRNAdb (95). We created wild-type and Δ*mod5* strains containing an additional tRNA^Tyr^GΨA gene under the control of a TetOn promoter and induced with doxycycline after confirming tRNA overexpression by northern blot to test for rescue of select phenotypes using droplet assays (**Fig. 6A, Fig. S9**).

**Figure 6.**
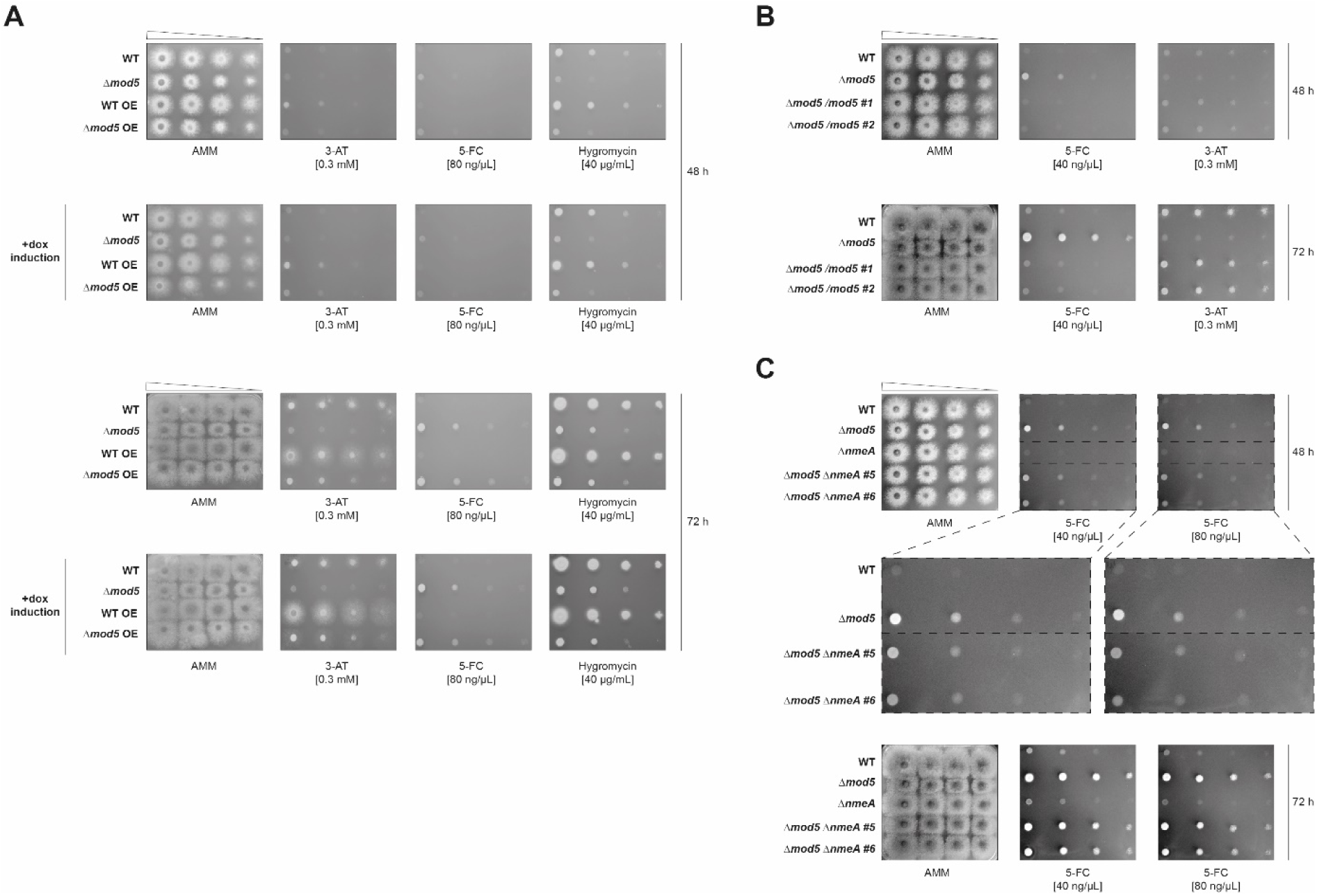
Deletion of *nmeA* but not tRNA^Tyr^GΨA overexpression reverts the 5-FC phenotype of Δ*mod5*. (A) The droplet growth assays for the wild-type and Δ*mod5* strain, as well as the respective tRNA^Tyr^GΨA overexpression strains, were grown for 48 h and 72 h. The strains were spotted and cultivated on AMM agar plates containing either 0.3 mM 3-AT, 40 µg/mL hygromycin, or 80 ng/µL 5-FC. The following displayed droplet growth assays were conducted with the wild type and Δ*mod5* and compared to either (B) two independent clones of reintroduced Δ*mod5/ mod5* or (C) Δ*nmeA* and, two independent clones of Δ*mod5* Δ*nmeA*. For (B) the strains were cultivated on AMM agar plates containing 40 ng/µL 5-FC or 0.3 mM 3-AT while for (C) the plates contained either 40 ng/µL or 80 ng/µL 5-FC. All droplet growth assays were incubated for 48 h and 72 h. Data are representative images from 3 biological replicates for (A-C).

Droplet assays on 3-AT supplemented AMM-agar plates (without doxycycline) recapitulated the Δ*mod5* phenotype, whereas the strain containing the tRNA-overexpression construct displayed a slight growth improvement (**Fig. 6A**, top panel). This could be explained by the well-known leakiness of the TetOn promoter (40). Induction of the promoter with doxycycline resulted in improved growth of Δ*mod5* on 3-AT (48 h and 72 h) in line with the proposed rescue effect by overexpression of hypomodified tRNA^Tyr^GΨA (**Fig. 6A**, +dox induction). Interestingly, the wild-type strain containing the same overexpression construct displayed similarly improved growth under normal conditions and after induction of hypomodified tRNA^Tyr^GΨA, supportive of a general role for tRNA^Tyr^GΨA in 3-AT stress response. Droplet assays with either 5-FC or hygromycin resulted in varying phenotypes. Contrasting the findings with 3-AT, overexpression of tRNA^Tyr^GΨA in Δ*mod5* did not revert the resistance phenotype of either stress. Notably, the wild type displayed a partial rescue on hygromycin upon tRNA overexpression (**Fig. 6A**). These findings imply a more complicated interplay of tRNA overexpression, translation, and stress response in the Δ*mod5* strain. Since a higher-than-normal dose of tRNA^Tyr^GΨA resulted only in a partial rescue of the modification deficient strain on the tested stressors it might be assumed that multiple tRNAs (e.g., cy-tRNA^Cys^GCA) are functionally affected by lack of i^6^A, necessitating a more challenging, combined tRNA-overexpression approach or alternatively higher overexpression driven by the more appropriate RNA Polymerase III. In line with this idea, complementing for the lack of *mod5* and correspondingly i^6^A37 resulted in a complete reversal of the 3-AT and 5-FC phenotypes (**Fig. 6B, Fig. S10A**), indicating the necessity for tRNA isopentenylation to cope with the stressors and most probably its presence on multiple isoacceptors.

Since our transcriptional data indicated a CPC-driven induction of MFS exporter *nmeA* (and two orthologs), we explored its importance for the 5-FC resistance of the *mod5* knockout. Correspondingly, we deleted *nmeA* in wild type and Δ*mod5* and repeated the droplet assays on AMM plates containing 5-FC (**Fig. 6C; Fig. S10B**) and separately overexpressed *nmeA* to demonstrate sufficiency in providing 5-FC resistance in *A. fumigatus* (**Fig. S11**). Knocking out *nmeA* resulted in no obvious phenotype compared to wild type but a notable increase in 5-FC susceptibility for the Δ*mod5* Δ*nmeA* strain. After 48 h, the double knockout displayed an increased sensitivity to 5-FC compared to Δ*mod5* but was still more resistant to the antifungal than wild type or Δ*nmeA* (**Fig. 6C**). These Δ*mod5-*to-Δ*mod5* Δ*nmeA* differences were milder after 72 h and with resistance potentially compensated by other NmeA orthologous exporters (e.g. *AFUB_043340* and/or *AFUB_028350* (**Fig. 5C**)) during longer growth periods. In sum, this data supports a role for NmeA as a main driver of 5-FC resistance in *A. fumigatus* Δ*mod5*.

## DISCUSSION

Antifungal resistance poses a major threat to the human population and puts at risk our already limited repertoire of antifungal drugs. Our understanding of antifungal drug resistance mechanisms centers primarily on genetic mutations, with limited examples of more transient forms of resistance. Here, we describe a mechanism of 5-FC/5-FOA drug resistance mediated by recognition of hypomodified tRNAs in the mold, *A. fumigatus* (**Fig. 7**). Loss, or potentially even downregulation, of Mod5 activity primes the cellular environment for enhanced resistance to 5-FC and 5-FOA by activation and upregulation of *cpcA* and CpcA-client gene *nmeA*, a known MFS transporter of nucleobases linked to 5-FC resistance in *A. nidulans* (*91*).

**Figure 7.**
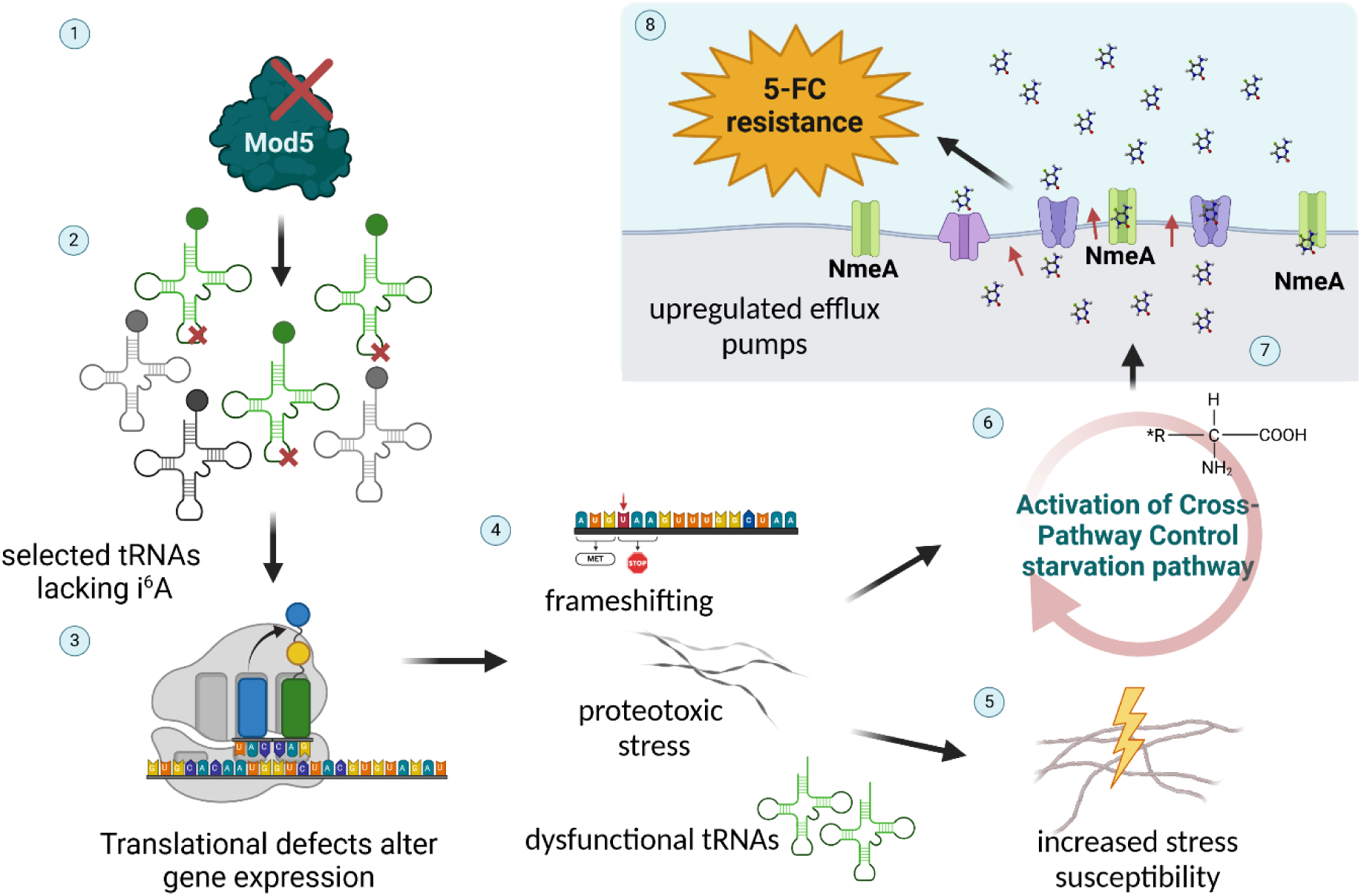
Proposed model of the 5-FC antifungal resistance mechanism of *A. fumigatus* Δ*mod5*. 1) Ablation of the tRNA-isopentenyl transferase Mod5 results 2) in the loss of i^6^A on specific target tRNAs. 3) As a result, gene expression is altered due to translational defects caused by 4) dysfunctional, hypomodified tRNAs, which likely leads to increased frameshifting, proteotoxic stress, and other consequences. 5) These defects manifest phenotypically as increased stress susceptibility in the *mod5*-knockout. 6) In parallel, the CPC system is prematurely activated, resulting in upregulation of various client genes including the nucleobase-specific NmeA efflux pump. 8) Higher levels of NmeA result in increased nucleobase efflux and 5-FC resistance. Created in BioRender. Blango, M. (2024) BioRender.com/k21b405

tRNA modifying enzymes, including Mod5, have mostly been investigated in non-pathogenic fungi like *S. cerevisiae* or *S. pombe* (16), with limited consideration of antifungal drug susceptibility or resistance. *S. cerevisiae* Δ*mod5* was shown to exhibit increased susceptibility to 5-FU, a derivative of the antifungal drug 5-FC that results in production of toxic derivatives (e.g., 5FUMP, 5FUTP), and an increased azole-resistance through formation of Mod5- amyloids and subsequent ablation of i^6^A (37,72). Together with studies showing the importance of IPTases in bacterial pathogens, we hypothesized that Mod5 could serve as a hub to coordinate responses to environmental stresses, including antifungal drugs in the stress-tolerant human fungal pathogen, *A. fumigatus* (21,22).

As expected, the *A. fumigatus mod5* knockout displayed sensitivity to agents like 3-AT and hygromycin, similar to many other anticodon stem-loop-tRNA modification deficient mutants (16,96). Surprisingly, the knockout grew on agar plates supplemented with 5-FC and 5-FOA, with a roughly 5-fold higher MIC than the wild type (**Fig. 1B**, **Table S6**). 5-FC has a documented low activity on *A. fumigatus* at neutral pH, where the fungus uses PacC and the CCAAT binding complex (CBC) to suppress the expression of purine cytosine transporter FcyB (79). This mechanism allows the fungus to effectively stop the uptake of 5-FC and is only reversed upon acidic environmental pH. Other forms of 5-FC resistance can be caused by mutation of downstream effectors of the 5-FC salvaging system including *fcyA*, *uprt*, *udpB*, and *urkA* (77,79,80). Interestingly, by performing a transcriptome and proteome analysis of the wild type and Δ*mod5* under non-stressed and 5-FC-stressed conditions, we were unable to find deregulation of the five 5-FC metabolizing genes that explained resistance. The only exception was a slight downregulation of *urkA* in the Δ*mod5* strain under normal growth conditions that was lost upon 5-FC treatment (**Fig. 2C**; **Table S8**). The *urkA* gene encodes for an enzyme that converts 5-fluorouridine to 5-fluorouridine monophosphate (5-FUMP); however, deletion of *urkA* only confers slight resistance (80), suggesting that deregulation of the salvage pathway factors is not responsible for the Δ*mod5* phenotype.

A closer interrogation of the transcriptional reprogramming induced upon 5-FC treatment of the wild type and knockout indicated similar pronounced adaptive stress responses. This included the prevention of DNA damage and translational impedance caused by 5-FC/5-FU, alongside a deregulation of various transmembrane transporters consistent with observations from other non-pathogenic and pathogenic fungi (**Fig. 2B, Table S8,12-14**; (77,81–83)). Transcriptomics revealed an induction of amino acid starvation responsive genes in Δ*mod5*, which was also partly observed in non-stressed conditions (**Fig. 2B**, **Table S8,14**). We identified these changes as part of cross-pathway control, which is regulated by the transcription factor CpcA to regulate over 500 genes (86,90). Quantification of transcriptionally autoregulated *cpcA,* as well as one client gene *argB,* confirmed premature activation of the CPC in the knockout, similar to Δ*mod5* in *S. cerevisiae* (70); but revealed substantial differences upon 3-AT and 5-FC treatment (**Fig. 4A,B**). Both agents led to CPC activation, but surprisingly *argB* mRNA levels were significantly higher in Δ*mod5* than in the wild type upon 5-FC stress. These results were surprising, since Δ*mod5* was highly susceptible to 3-AT unlike 5-FC, and we had hypothesized that this susceptibility was caused by additional activation of the prematurely induced CPC (70,76,96,97). This paradox was explained by the upregulation of CpcA-client gene *nmeA* (**Fig. 5A,B**). Overexpression of *A. nidulans nmeA* was shown to confer 5-FC resistance (91), which we confirmed in *A. fumigatus* (**Fig. S11**). Notably, the *Ani_nmeA* promoter region also contains a CpcA binding site but remains lowly expressed under several stress conditions contrasting the 5-FC-induced, CPC-driven *nmeA* regulation in

*A. fumigatus* (**Fig. 5A,B**) (91). Our experiments displayed the disproportionate CPC activation in Δ*mod5* in form of (i) higher expression of this and other *Ani_nmeA* orthologous exporters (**Fig. 5A-C**), while (ii) also showing a reduction of 5-FC-derivative 5-FU levels incorporated into tRNA transcripts (**Fig. 5D**). Interestingly, the decreased 5-FU incorporation did not result in significantly less perturbation of uridine-directed modification like Ψ and m^5^U in Δ*mod5* (**Fig. 5D**), although displaying a tendency that might increase over a longer 5-FC/5-FU treatment period. Deleting *nmeA* in the tRNA modification knockout ultimately reverted, although not completely, the 5-FC resistance phenotype (**Fig. 6C**). Due to the observed incomplete rescue of the double knockout after 48 h and 72 h incubation, we speculate that additional premature CPC-induced exporters e.g., *AFUB_043340* and/or *AFUB_028350* (**Fig. 5C**) might partially compensate for the lack of NmeA in the Δ*mod5* background.

We propose a model where loss of Mod5-dependent installation of i^6^A37 on specific tRNAs (16,35) contributes to the observed 5-FC resistance phenotype of the *mod5* deletion. Ablation of tRNA modifications in the anticodon stem-loop is known to influence translational capacity and proteome stability, resulting in the reprogramming of the transcriptome and activation of premature starvation response programs (70,76,96). In agreement, we observed an induction of starvation responses, specifically the CPC in Δ*mod5* (**Fig. 2B**, **4A,B**), which leads to the induction of *nmeA* (and orthologues) and 5-FC resistance (**Fig. 5A,B**; **Fig. 6C; Table S8**). One explanation might be offered by MoTTs that are enriched for i^6^A-dependent codons and decreased in translational efficiency in the knockout (88). In line with this hypothesis, our proteomic and codon-frequency analyses in non-stressed and 5-FC-stressed conditions displayed a negative correlation for i^6^A-dependent tyrosine and cysteine and a positive correlation for arginine codons with protein abundance in the knockout (**Fig. 3B-D**, **Fig. S7,8**). 5-FC treatment in the wild type resulted in decreased levels of i^6^A (**Fig. 1A**), which (i) indicates the influence of 5-FU even on non-uridine tRNA modifications and (ii) might hint at activation of the CPC-driven resistance axis upon longer periods of antifungal treatment. Of note, certain tRNA modifiers (*trm61* (*AFUB_069660*), *AFUB_015050* (*TYW4* in *S. cerevisiae*), *trm82* (*AFUB_005430*), and *AFUB_090820* (*QRI7* in *S. cerevisiae*)) were also among the decreased proteins in Δ*mod5*, implying a more complex, coordinated change of the epitranscriptome than initially expected (**Table S17**).

Many phenotypes observed for tRNA modification-deficient mutants can be rescued by providing an excess of the affected hypomodified tRNA(s) (26,76,98). Applying this strategy to Δ*mod5* transformed with an additional TetOn promoter-controlled tRNA^Tyr^_GΨA_ gene failed to completely rescue the phenotypes under different stressors (**Fig. 6A**). The tRNA overexpression on 3-AT rescued the knockout growth defect and provided an advantage to the wild type, while the phenotypes on hygromycin and 5-FC were unchanged. It is known from other tRNA modifier mutants that a combined tRNA overexpression is necessary in some cases to perform an efficient rescue (26). With the help of Nano-tRNAseq it was possible to identify four cytoplasmic Mod5 tRNA modification targets in *A. fumigatus* (**Fig. 1B, 3B, Fig. S3)** of which additional overexpression of specific isoacceptors like tRNA^Cys^GCA might be necessary to revert growth defects (16). In contrast to this rationale, there was no significant negative correlation observed between wild type and knockout under 5-FC stress (**Fig. S8**). Notably, the strategy performed here for tRNA overexpression seemed to only result in a mild excess of tRNA^Tyr^GΨA, which might also influence the rescue efficiency on the *mod5* knockout phenotypes (**Fig. S9**).

Of note, eukaryotic IPTases are also known to secondarily function as factors in tRNA gene-mediated silencing, which could potentially contribute to the Δ*mod5* 5-FC resistance phenotype (35). Performing a correlation analysis on tRNA gene-to-protein-coding gene distances using our transcriptome data revealed an overall closer proximity of induced genes to tRNAs in the knockout (**Fig. S12**). This correlation might hint at a contribution of this mechanism to the Δ*mod5* antifungal resistance reminiscent of epimutation-based mechanisms linked to antifungal resistance in other organisms, including fungal pathogens (99,100). Certainly, more experimental data is necessary to confirm such an involvement.

In conclusion, we provide data for an unexpected mechanism of *A. fumigatus* resistance against the antifungal drug 5-FC, occurring after deletion of the tRNA modifier Mod5. The corresponding ablation of i^6^A37 results in impeded translation and transcriptional reprogramming, which culminates in the CPC-driven activation of MFS exporter *nmeA* (and orthologs). According to the nature of this resistance mechanism, we postulate that conditions resulting in tRNA hypomodification or other forms of premature CPC activation may provide an unintended, transient benefit to *A. fumigatus* in coping with 5-FC and potentially other antifungal drugs, with clear implications for therapeutic success against this dangerous pathogen.

## DATA AVAILABILITY

The mass spectrometry proteomics data were deposited in the ProteomeXchange Consortium via the PRIDE partner repository (101) with the dataset identifier PXD048351. The mRNA-seq dataset can be found at GEO under identifier GSE251655. The Nano-tRNAseq dataset was also deposited at GEO with the identifier GSE276346.

## SUPPLEMENTARY DATA

Supplementary Data are available at NAR online.

## ACKNOWLEDGEMENTS

We would like to thank both Axel Brakhage and Amelia Barber for helpful discussions of this manuscript. We would also like to thank Bhawana Israni, Lukas Schrettenbrunner, and Nathalie Seiler for experimental support and excellent technical assistance.

## FUNDING

The work presented here was generously supported by the Federal Ministry for Education and Research (BMBF: https://www.bmbf.de/), Germany, Project FKZ 01K12012 “RFIN – RNA-Biologie von Pilzinfektionen”. Additional funding support came from the Deutsche Forschungsgemeinschaft (DFG; German Research Foundation) under Germanýs Excellence Strategy – EXC 2051 – Project-ID 390713860, the DFG-funded Collaborative Research Center/Transregio FungiNet 124 ‘Pathogenic fungi and their human host: Networks of Interaction’ (210879364, project INF and Z2), and the BMBF-funded “PerMiCCion” project (Project ID 01KD2101D).

## CONFLICT OF INTEREST

The authors declare no conflict of interest.

## Notes

### Competing Interest Statement

The authors have declared no competing interest.

### Summary of Updates

The manuscript was updated to more adequately explain the mechanism of 5-FC resistance observed. New data was added for tRNA modification status in the knockout, Mod5 target tRNAs by Nano-tRNAseq, and rescue of the observed antifungal resistance phenotypes.

