## Supplemental Material for "tRNA hypomodification facilitates 5-fluorocytosine resistance via cross-pathway control system activation in *Aspergillus fumigatus*"

† Joint Authors are listed in alphabetical order.

**SUPPLEMENTAL TABLES**

Table S1. *Aspergillus fumigatus* strains used in this study. Annotated are the name of the strain, the genotype and originating source of the strain.

| Strain | Genotype | Reference |
|---|---|---|
| **CEA17 Δ*akuB*** $^{KU80}$ | *akuB* $^{KU80}$*::pyrG*; PyrG+, Δ*akuB* $^{KU80}$ | (1) |
| **Δ*mod5*** | CEA17 Δ*akuB* $^{KU80}$, *AFUB_093210::ptrA* | This study |
| **Δ*mod5/mod5*** | Δ*mod5*, *pyrG::AFUB_093210, ble* | This study |
| **Δ*nmeA*** | CEA17 Δ*akuB* $^{KU80}$, *AFUB_005530::hygR* | This study |
| **Δ*mod5*Δ*nmeA*** | Δ*mod5*, *AFUB_005530::hygR* | This study |
| **CEA17 Δ*akuB*** $^{KU80}$ **tRNA** $^{Tyr}_{GUA}$ **OE** | CEA17 Δ*akuB* $^{KU80}$, *pyrG::TetON_AFUB_013600, ble* | This study |
| **Δ*mod5* tRNA** $^{Tyr}_{GUA}$ **OE** | CEA17 Δ*akuB* $^{KU80}$ Δ*mod5, pyrG::TetON_AFUB_013600, ble* | This study |
| **Δ*cntA::nmeA*** $^{PxylP}$ | CEA17 Δ*akuB* $^{KU80}$, *cntA::PxylP_AFUB_005530_TAtTrpC* | This study |

Table S2. Oligonucleotides used in this study.

| Name | 5'-3' sequence | Target gene | Reference/ source |
|---|---|---|---|
| **Oligonucleotide primers** | | | |
| mod5-KO-o1 | ACCTCCAATGAGCAATGTCC | *AFUB_093210*, 5'flank, F | This study |
| mod5-KO-o2 | GGCCTGAGTGGCCATCGAATTC GGTTGTTTCGTTACGAGTCC | AFUB_093210 5' flank, R | This study |
| mod5-KO-o3 | GAGGCCATCTAGGCCATCAAGC ATACATAGTGACTCGAGCCG | *AFUB_093210*, 3'flank, F | This study |

| | | | |
|---|---|---|---|
| mod5-KO-o4 | AAGCTCCTAAAGTTCTGGGC | *AFUB_093210*, 3'flank, R | This study |
| mod5-KO-o5 | GAATTCGATGGCCACTCAGGCC | *ptrA,*F | This study |
| mod5-KO-o6 | ATGGCCTAGATGGCCTCTTGC | *ptrA,* R | This study |
| CpcA-o1 | CTTGGATGCTGCAATCGCTC | *AFUB_069420*, *cpcA*, F | This study |
| CpcA-o2 | TGGAGTGCTTAACACCGGAC | *AFUB_069420*, *cpcA*, R | This study |
| CoxV-o1 | ATCTGTTCGCCAAGCCCAAG | *AFUB_058180*, *coxV*, F | (2) |
| CoxV-o1 | TCACTGCTGACACCGTAGAG | *AFUB_058180*, *coxV*, R | (2) |
| ArgB-o1 | CGTCAACGTGTCTCCCTCAA | *AFUB_064280*, *argB*, F | This study |
| ArgB-o2 | GGTAGCGAACTCGGTAGGTG | *AFUB_064280*, *argB*, R | This study |
| pyrG-o1 | GAGCTACTCAGACATAGCTAC | *Afu2g08360*, 5'flank, F | This study |
| pyrG-o2 | CGTGGGAATGGAGGGTTATA | *Afu2g08360*, 5'flank, R | This study |
| $tRNA^{Tyr}_{G\Psi A}$-OE-o1 | AGACTTAGGCCGCTATGTACGACCTGGG | *AFUB_013600,* $tRNA^{Tyr}_{G\Psi A}$, F | This study |
| $tRNA^{Tyr}_{G\Psi A}$-OE-o2 | GGCATAAATCGAATGTCCGCAAAAGTTGTCCTAATCCCC | *AFUB_013600,* $tRNA^{Tyr}_{G\Psi A}$, R | This study |
| $tRNA^{Tyr}_{G\Psi A}$-OE-o3 | GCGGACATTCGATTTATGCC | *A. nidulans* Ttef, F | This study |
| $tRNA^{Tyr}_{G\Psi A}$-OE-o4 | TTGGGATGAATTTTGTATGC | *A. nidulans* Ttef, R | This study |

| | | | |
|---|---|---|---|
| $tRNA^{Tyr}_{G\Psi A}$-OE-o5 | GCATACAAAATTCATCCCAATTTCCTCTTGGAGCAAAAGT | *Afu2g08360* flanks_*Ttef*, F | This study |
| $tRNA^{Tyr}_{G\Psi A}$-OE-o6 | GAATTCGAGCTCGGTACCCGAATTCGGGACTTGGGGCCTATTCAT | *Afu2g08360* flanks_*pUC18_PyrG*, R | This study |
| $tRNA^{Tyr}_{G\Psi A}$-OE-o7 | TCGACTCTAGAGGATCCCCGAATTCGAGCTACTCAGACATAGCTA | *Afu2g08360* flanks_*pUC18_PyrG*, F | This study |
| $tRNA^{Tyr}_{G\Psi A}$-OE-o8 | GTACATAGCGGCCTAAGTCTTGTGATGTGATGGAGTTGAG | *AFUB_013600_TetON_ble*, R | This study |
| Mod5-compl-o1 | TACGTATCTAGAACTAGTCTCGACTCATCTAGGCTGGAGGGGA | *AFUB_093210*, CDS -1 kb, F | This study |
| Mod5-compl-o2 | TTGGGATGAATTTTGTATGCATCAGGAACGACCGGGAC | *AFUB_093210*, CDS + 1 kb, R | This study |
| Mod5-compl-o3 | GCATACAAAATTCATCCCAATTTCCTCTTGGAGCAAAAGT | *Afu2g08360*, A. niduland Ttef, F | This study |
| Mod5-compl-o4 | AGACTAGTTCTAGATACGTATCTAGAAAGAAGGATTACCT | *TetON_ble*, *AFUB_093210_R* | This study |
| Mod5-compl-o5 | CTGCGGCGTTGAAGGCTGCT | *AFUB_093210_CDS_R* | This study |
| NmeA-o1 | TCCCCTTGCAAGTGAAGTCT | *AFUB_005530*, 5‘flank, F | This study |
| NmeA-o2 | TCTCCCTCAACCTCACCAAC | *AFUB_005530*, 3‘flank, F | This study |
| NmeA-o3 | CGGAAGCAATTGGACTTCTG | *AFUB_005530*, *hph* | This study |
| NmeA-KO-o1 | CCGGCTCGGTAACAGAACTAACGGCGTAACCAAAAGTCAC | *PgpdA-hph* cassette from *pAN7.1* | This study |

| | | | |
|---|---|---|---|
| NmeA-KO-o2 | GGGAGCATATCGTTCAGAGCTCTTGA CGACCGTTGATCTG | *PgpdA-hph* cassette from *pAN7.1* | This study |
| NmeA-KO-o3 | AGTGAATTCGAGCTCGGTACTCCCCT TGCAAGTGAAGTCT | *AFUB_005530*, 5‘flank, F | This study |
| NmeA-KO-o4 | TAGTTCTGTTACCGAGCCGGCCCCAC TGCCTGTCTTCTA | *AFUB_005530*, 5‘flank, R | This study |
| NmeA-KO-o5 | GCTCTGAACGATATGCTCCCCGAAGC GTTTGTGGCTTATT | *AFUB_005530*, 3‘flank, F | This study |
| NmeA-KO-o6 | TTACGCCAAGCTTGCATGCCTCTCCC TCAACCTCACCAAC | *AFUB_005530*, 3‘flank, R | This study |
| NmeA-OE-o1 | TTCTTCAAAGAAGGCGTTTTGGACAC CCTGAAAGGTCGTCGCGTTCTTAACT GATGCGAGCAACAGTATGC | 5‘*AfucntA*-*PcPxylP*-fw | This study |
| NmeA-OE-o2 | AATTCGCAAAATGAATATGATAAAAAT ACAGTTCTTGTGATTTGCAGGCAAGG GTTGAGTACGAGATTGGGG | 3‘*AfucntA_AtTrpC*-rv | This study |
| NmeA-OE-o3 (pX-FW.2) | CCATGGCAGCAGTGATTTCA | pΔ*fcyB*_mKate2 *xyl* | (3) |
| NmeA-OE-o4 (pX-RV.2) | GGTTGGTTCTTCGAGTCGATG | pΔ*fcyB*_mKate2 *xyl* | (3) |
| NmeA-OE-o5 | ATCGACTCGAAGAACCAACCATGAGG GCACATGCGGG | *PxylP_ AFUB_005530*-F | This study |
| NmeA-OE-o6 | TGAAATCACTGCTGCCATGGTCATGC TCTGACAAGTAACCG | *AttrpC_ AFUB_005530*-R | This study |
| NmeA-OE-o7 | ACTGGGGCTTTTTCTGGACT | 5’*AfucntA*-F | This study |
| NmeA-OE-o8 | GGTTTGGATATGCCCTTGAG | 5’*AfucntA*-R | This study |

| | | | |
|---|---|---|---|
| **Oligonucleotide probes – Northern blot** | | | |
| tRNA$^{Tyr}_{G\Psi A}$-p1 | CCCGAGCCGGAATCGAGCCG | cy-tRNA-Tyr BP | This study |
| **Oligonucleotide probes – Southern blot** | | | |
| *mod5* knockout confirmation | Mod5-o3 + Mod5-o4 | *AFUB_093210*, 3'flank | This study |
| tRNA$^{Tyr}_{G\Psi A}$ confirmation | pyrG-o1 + pyrG-o2 | *AFUB_013600,* tRNA$^{Tyr}_{G\Psi A}$, 5'flank | This study |
| *nmeA* knockout confirmation | NmeA-o1 + NmeA-o3 | *AFUB_005530*, 5'flank | This study |
| *nmeA* overexpression confirmation | NmeA-OE-o7 + NmeA-OE-o8 | *cntA*, 5'flank | This study |

**SUPPLEMENTAL FIGURES**

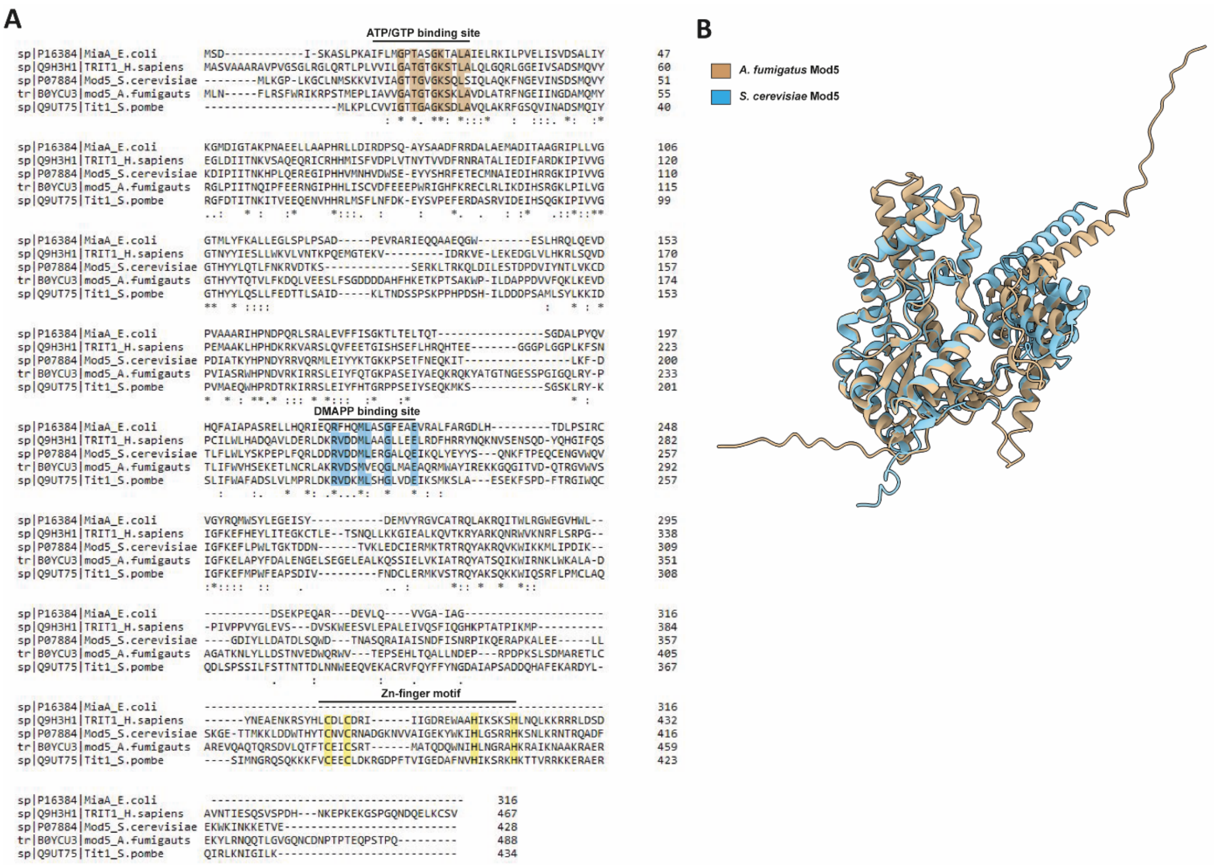


Figure S1. *Aspergillus fumigatus* Mod5 (*AFUB_093210*) shares amino acid similarity and domain conservation with IPTases from other organisms. (**A**) The amino acid sequences of *A. fumigatus* Mod5 and orthologs from *E. coli* (MiaA), *Homo sapiens* (TRIT1), *S. cerevisiae* (Mod5), and *S. pombe* (Tit1) were retrieved from the UniProt website and compared with the help of the Clustal Omega online tool (4-6). Conserved domains (**ATP/GTP binding site**, **DMAPP binding site**, **Zn-finger motif**) and associated amino acids (brown, blue and yellow) are indicated. "*" = conserved amino acid, "." = conservation between groups with weak similarity, ":" = conservation between groups of strongly similar properties. (**B**) Overlay of *S. cerevisiae* and *A. fumigatus* Mod5 structures. Predicted Alphafold structures were downloaded from https://www.uniprot.org/ and superimposed using the ChimeraX software's Matchmaker tool.

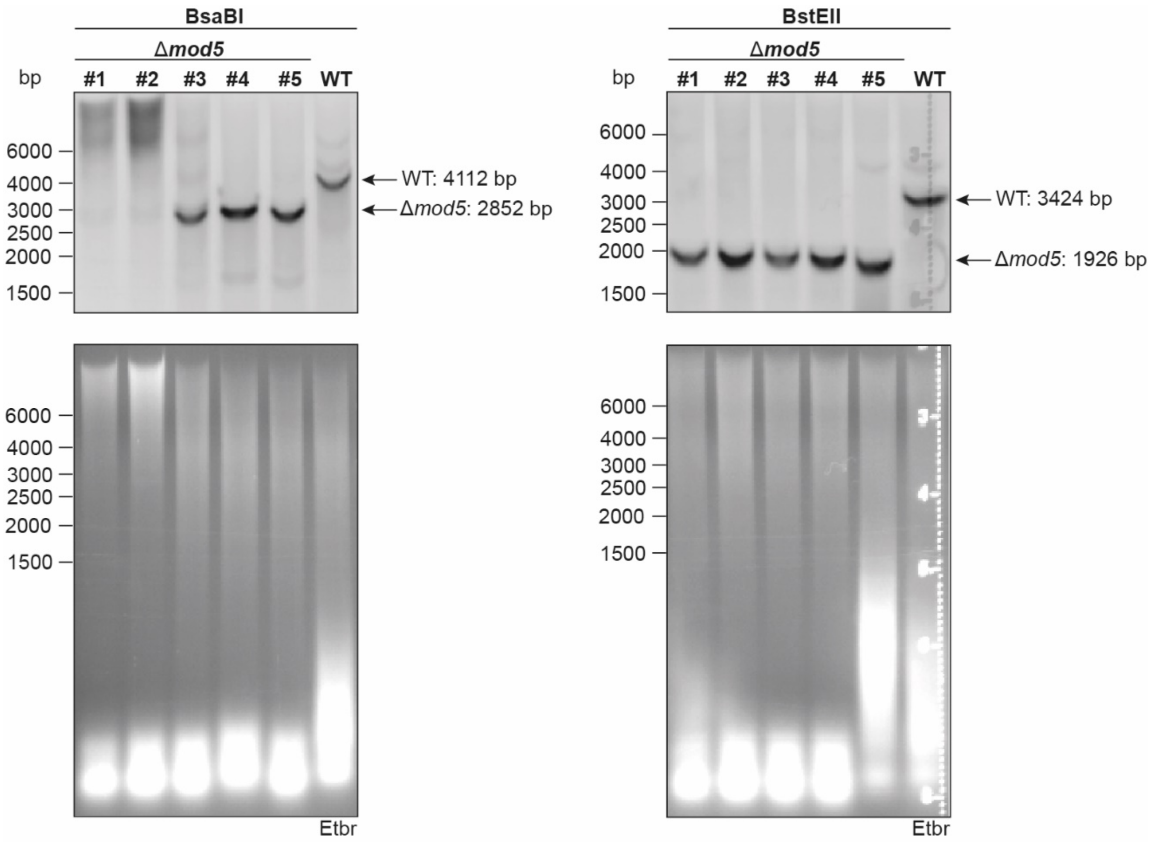


Figure S2. Confirmatory Southern blot to validate successful deletion of *mod5*. The WT or transformation clones were subjected to DNA isolation and 10 µg of the isolates were digested overnight with either BsaBI or BstEII. A successful insertion is represented by a digestion product of either 2,852 bp (left panel) or 1,926 bp (right panel), a negative result with a band at 4,112 bp and 3,424 bp, respectively. Confirmation of digestion is shown below with ethidium bromide (Etbr) stained agarose gel.

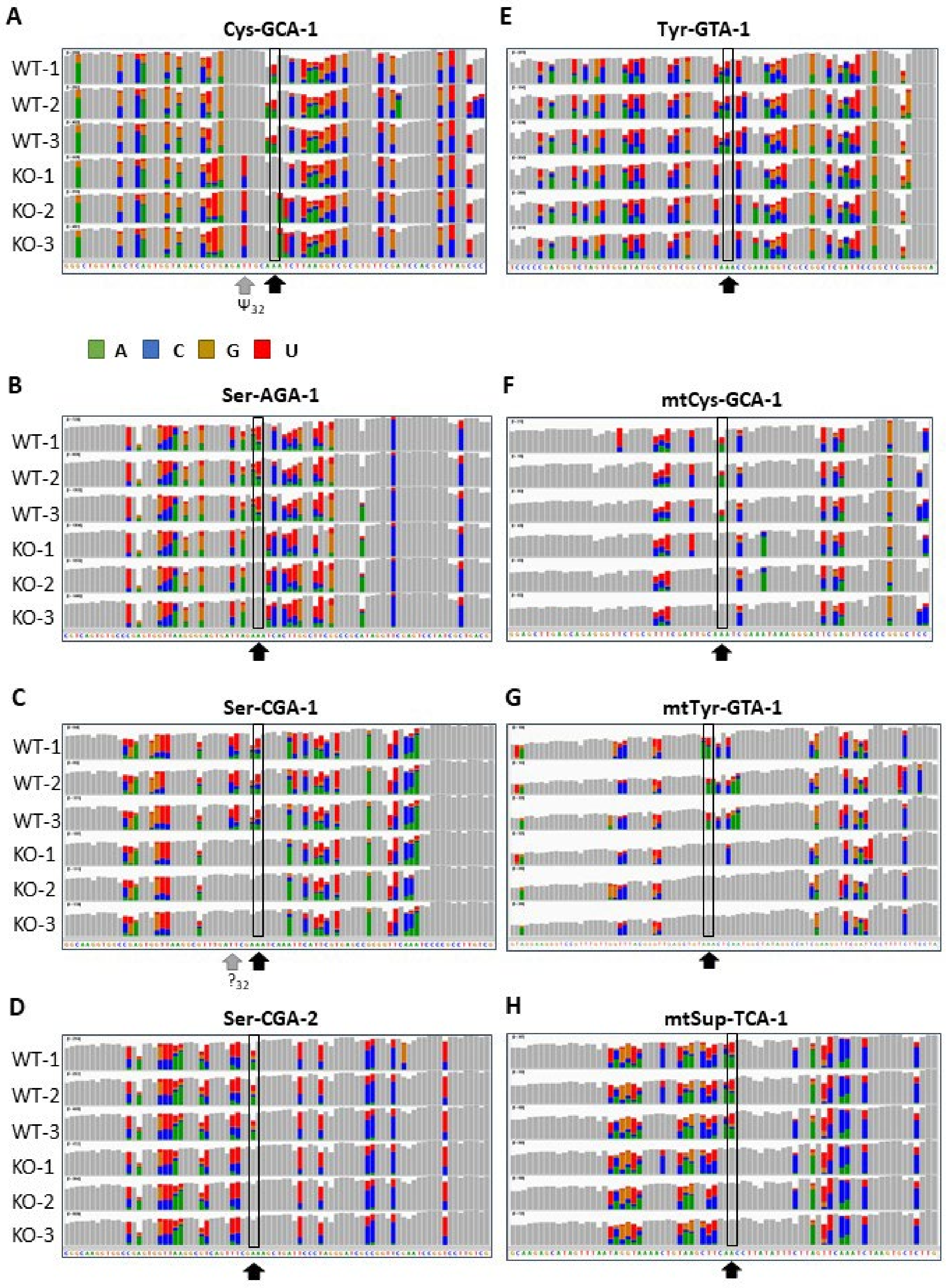


Figure S3. Eight different cytoplasmic and mitochondrial (mt) Mod5 tRNA modification targets were detected by Nano-tRNAseq. Displayed are Integrated Genome Viewer (IGV) snapshots of the indicated tRNAs and their sequences with grey (mismatch frequency <0.2) and coloured bars (mismatch frequency >0.2). Mod5-dependent isopentenylation of position 37 adenosines and its absence in the respective knockout is indicated by a black arrow and box. The grey arrows indicate the detection of additional mismatch (i.e., modification) variations in

$tRNA^{Ser}_{CGA}$-1 and $tRNA^{Cys}_{GCA}$ together with the presumed modifications at this position according to the MODOMICS database (7).

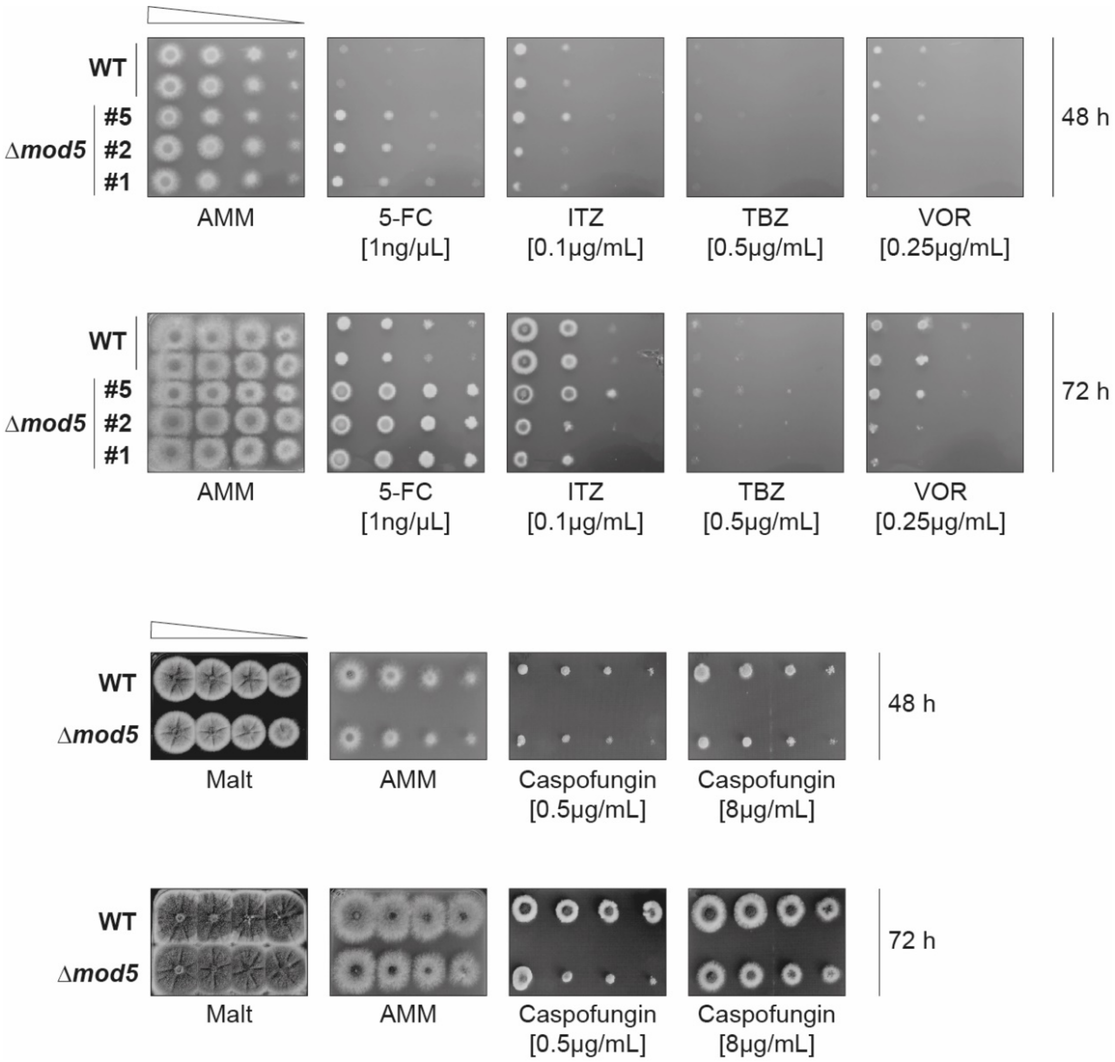


Figure S4. The deletion of *mod5* results in varying phenotypes against the antifungals caspofungin and multiple azoles. Strains were spotted and cultivated on AMM agar plates containing 0.5 µg/mL and 8 µg/mL caspofungin, 1 ng/µL 5-FC, 0.1 µg/mL itraconazole (ITZ), 0.5 µg/mL tebuconazole (TBZ) or 0.25 µg/mL voriconazole (VOR). WT and Δ*mod5* grown on AMM and Malt agar plates served as controls. Documentation of the different assays was carried out after 48 h and 72 h. Data are representative images from three biological replicates.

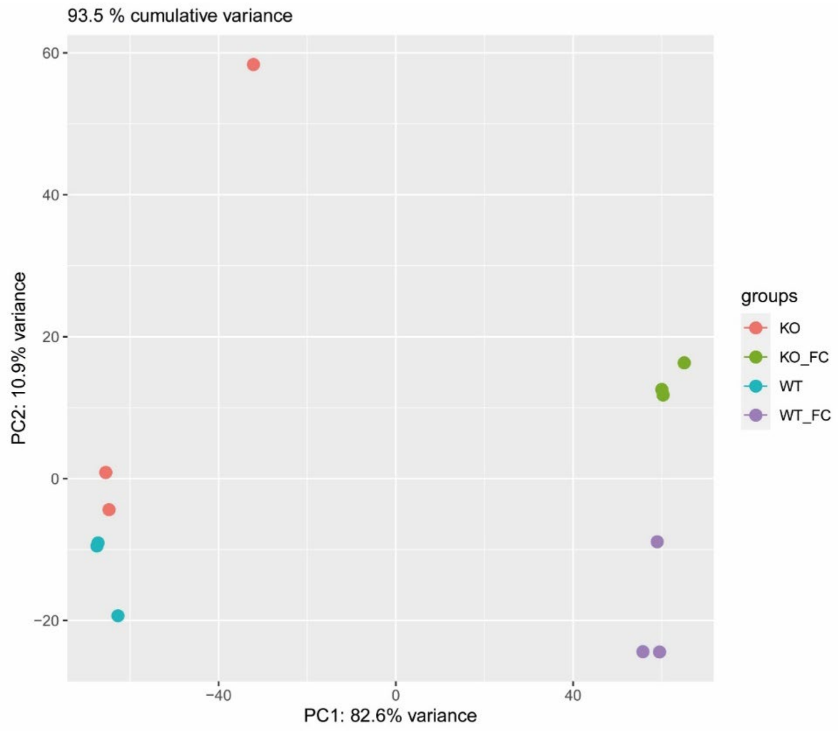


Figure S5. Principal component analysis of the transcriptome data reveals a clear separation of non-stressed and 5-FC stressed samples. The biological triplicates of both strains and conditions are represented by differently coloured dots. 93.5 % variability among the different sample groups is established with PC1 and PC2.

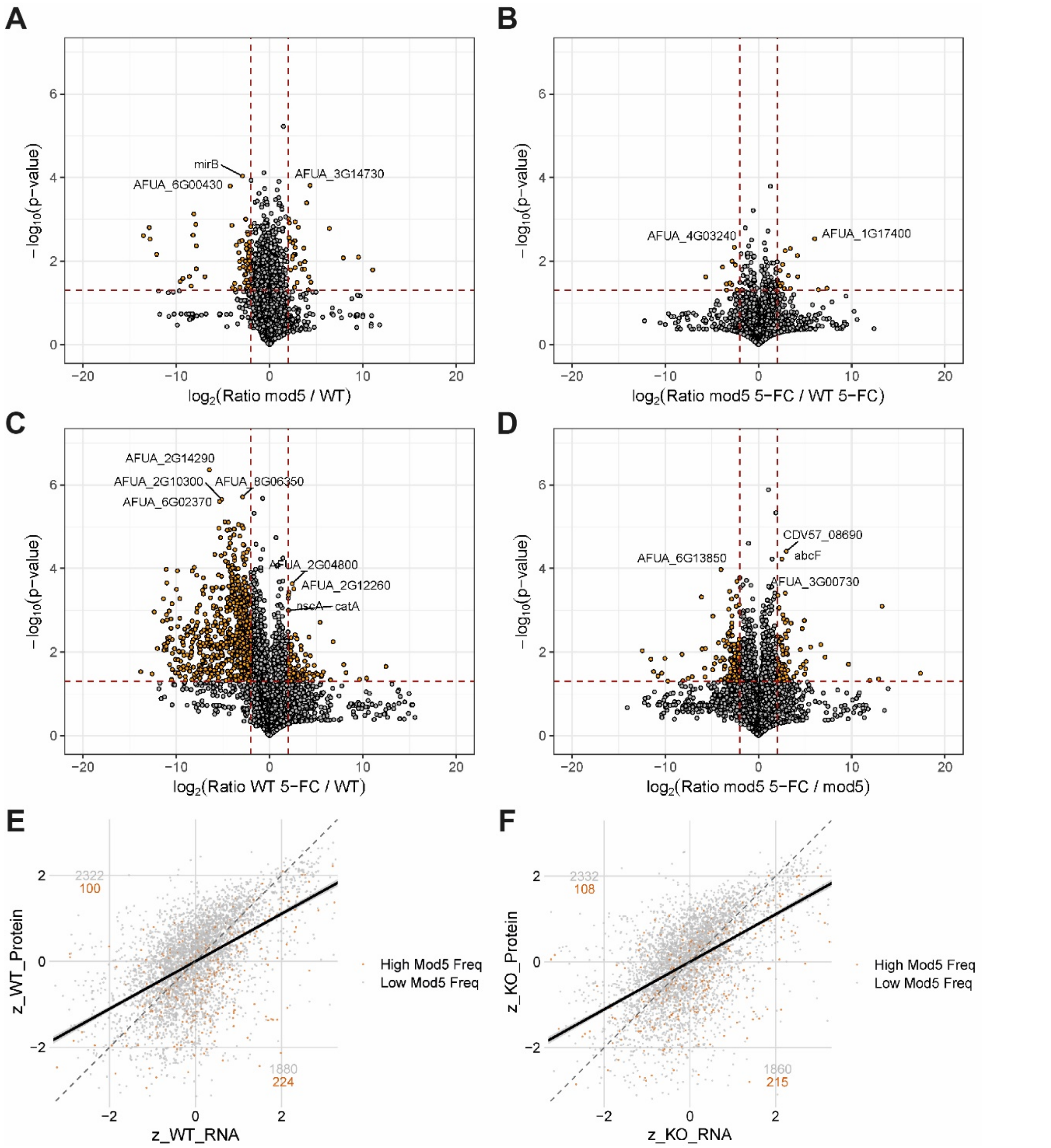


Figure S6. Proteomic changes correlate with transcriptome variations upon deletion of *mod5* or 5-FC treatment. (**A-D**) Volcano plots of the different comparisons display changes of the Δ*mod5* and WT proteome under non-stressed and 5-FC stressed conditions. The -$\log_{10}$ transformed *p*-values are given on the y-axis while the x-axis displays the $\log_2$FC. The dots in orange represent significantly (*p*-value≤0.05) changed protein abundance with a $\log_2$FC≥2. In each case, proteins were only included in the comparison if there were quantified in at least two biological replicates of one of the conditions in the proteomics. (**E-F**) Z-scored protein abundances and transcriptome data of the WT (**E**) and Δ*mod5* (**F**) were correlated and plotted with the help of the R package ggplot2 version 3.4.4. Only proteins that were quantified in at least two biological replicates were included. Orange dots correspond to protein-coding genes with Mod5 target codon usage more than one standard deviation above the mean.

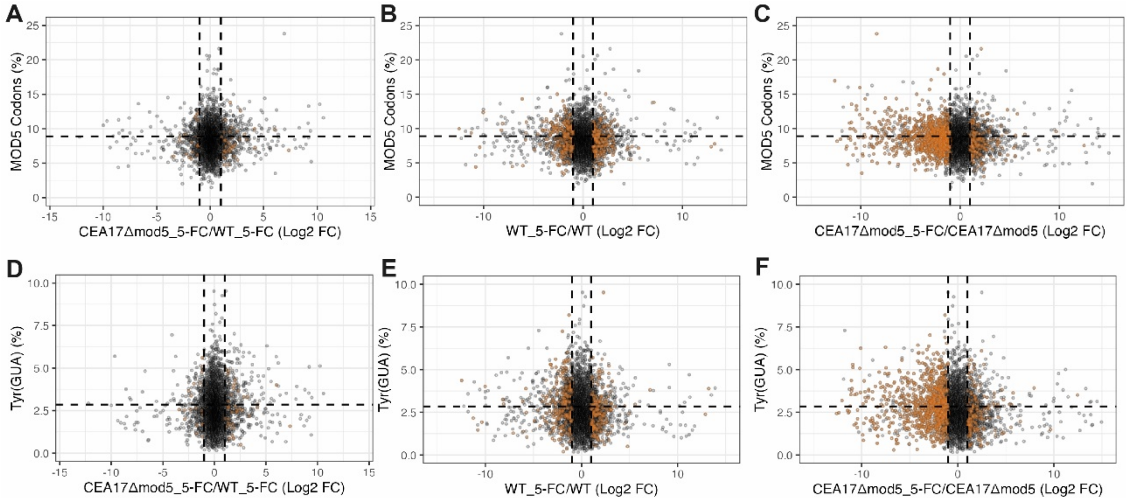


Figure S7. Proteins with decreased abundance in the *mod5* knockout display a higher tyrosine frequency. Plots display relative protein levels ($log_2FC \geq 2$) of the different comparisons conducted for the indicated strains upon different conditions (+/- 5-FC) versus all Mod5-dependent codons (**A-C**) or tyrosine frequency (**D-F**) per open-reading frame (**Table S16-20**). Orange dots indicate an enrichment for Mod5-dependent codons/tyrosine frequency in the respective protein.

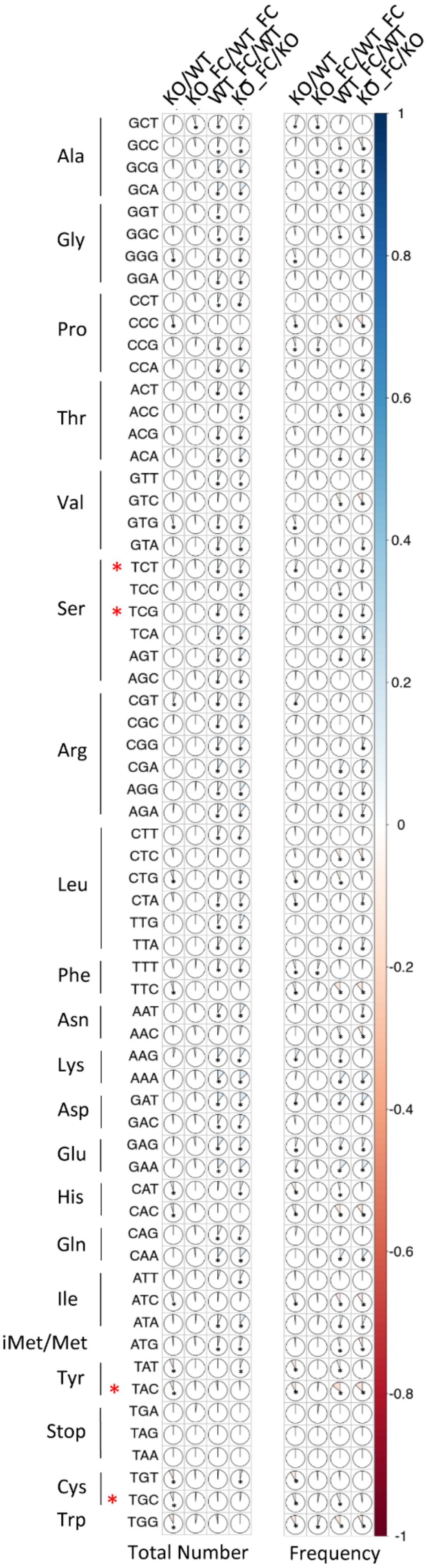

KO/WT
KO_FC/WT_FC
WT_FC/WT
KO_FC/KO
Ala
Gly
Pro
Thr
Val
Ser
Arg
Leu
Phe
Asn
Lys
Asp
Glu
His
Gln
Ile
iMet/Met
Tyr
Stop
Cys
Trp
GCT GCC GCG GCA GGT GGC GGG GGA CCT CCC CCG CCA ACT ACC ACG ACA GTT GTC GTG GTA TCT TCC TCG TCA AGT AGC CGT CGC CGG CGA AGG AGA CTT CTC CTG CTA TTG TTA TTT TTC AAT AAC AAG AAA GAT GAC GAG GAA CAT CAC CAG CAA ATT ATC ATA ATG TAT TAC TGA TAG TAA TGT TGC TGG
Total Number
Frequency
1
0.8
0.6
0.4
0.2
0
-0.2
-0.4
-0.6
-0.8
-1

Figure S8. Correlation between select Mod5 codon usage and protein abundance in the *mod5* deletion. Spearman's rank correlation coefficients were determined for the number or frequency of each triplet codon compared to the $Log_2FC$ values determined in the proteomics analysis (**Table S16**). Only proteins quantified in at least two biological replicates for one of the samples in a particular comparison were included. * = *A. fumigatus* Mod5-affected codons.

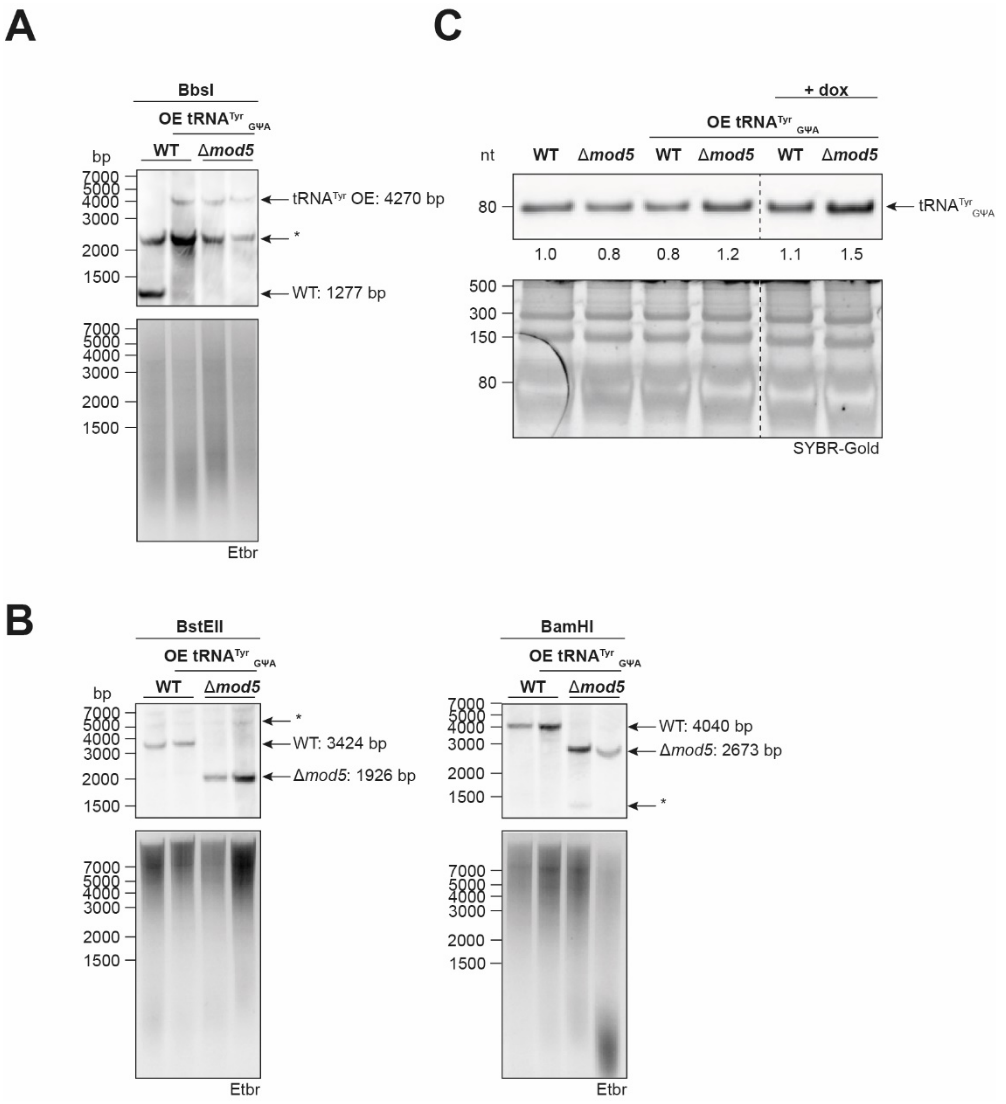


Figure S9. Confirmation of the tRNA overexpression strain construction. Representative Southern blots confirming proper construction of the (**A**) tRNA overexpression strain and deletion of (**B**) *mod5*. Successful insertion resulted in a digestion product of either 4,270 bp (BbsI, **A**), 1,926 bp (BstEII, **B**, left panel) or 2,673 bp (BamHI, **B**, right panel) while a negative result is represented with a band at 1,277 bp (**A**), 3,424 bp (**B**, left panel) and 4,040 bp (**B**, right panel). *=unspecific bands. (**C**) Northern blot to confirm doxycycline induced overexpression of $tRNA^{Tyr}_{GUA}$ of the indicated strains. Calculated arbitrary values were measured with the help of ImageJ. The blot is representative for three independent replicates.

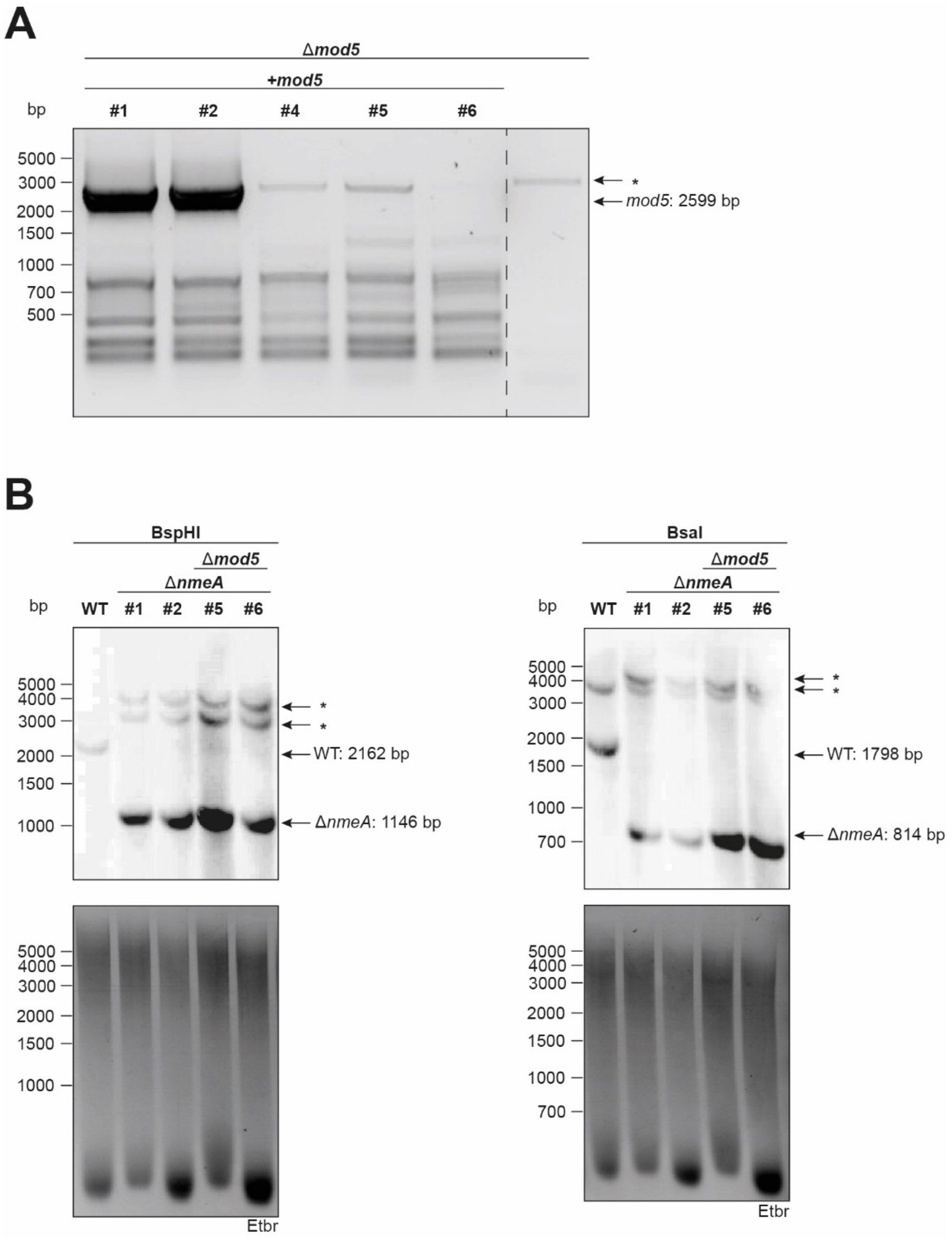


Figure S10. Genetic Confirmation of the *nmeA* knockout and Δ*mod5* complementation. (**A**) Complementation of Δ*mod5* with a reintroduced *mod5* gene was confirmed via PCR on the isolated DNA of the indicated strains. Successful reintroduction resulted in a PCR-product of 2599 bp using the primers Mod5-compl-o1 and Mod5-compl-o2 (**Table S2**). (**B**) DNA of the WT or the tested transformation clones were subjected to overnight digestion with either BspHI or BsaI and afterwards utilized for confirmatory Southern blots. Successful insertion is represented by a digestion product of either 1,146 bp (left panel) or 814 bp (right panel) while a negative result is indicated with a band at 2,162 bp and 1,798 bp, respectively. *=unspecific bands.

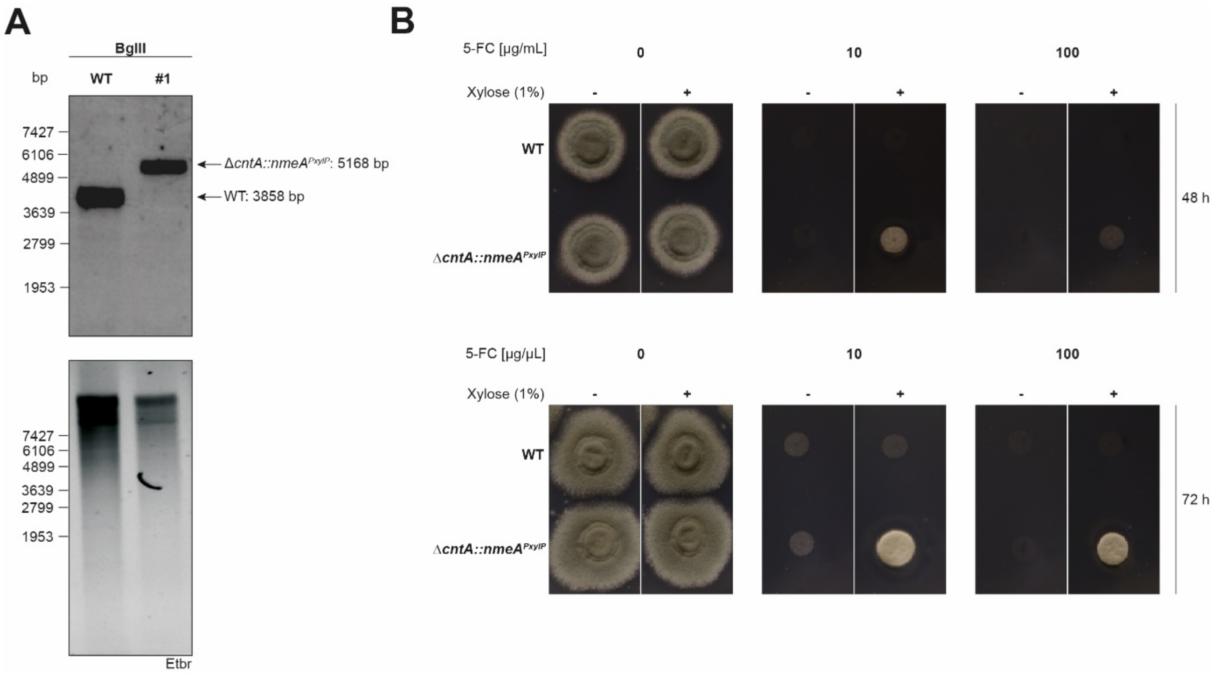


Figure S11. Overexpression of *nmeA* results in 5-FC resistance. (**A**) A xylose inducible *nmeA* overexpressing strain was constructed, resulting in *PxylP-nmeA* and was introduced in the counter-selectable marker locus *cntA* (8). Successful transformation was confirmed by digestion of the wild type (WT) and transformant genomic DNA with BglII overnight and performing a southern blot. Construct insertion is represented with a digestion product of 5168 bp while a negative result is indicated with a band at 3858 bp. (**B**) Strains were spotted and cultivated on pH 5-adjusted AMM agar plates containing 10-100 µg/mL 5-FC with/without 1% xylose. WT and Δ*cntA::nmeA*$^{PxylP}$ grown on AMM agar plates (pH5, +/- 1% xylose) served as controls. Documentation of the different assays was carried out after 48 h and 72 h. Data are representative images from three biological replicates.

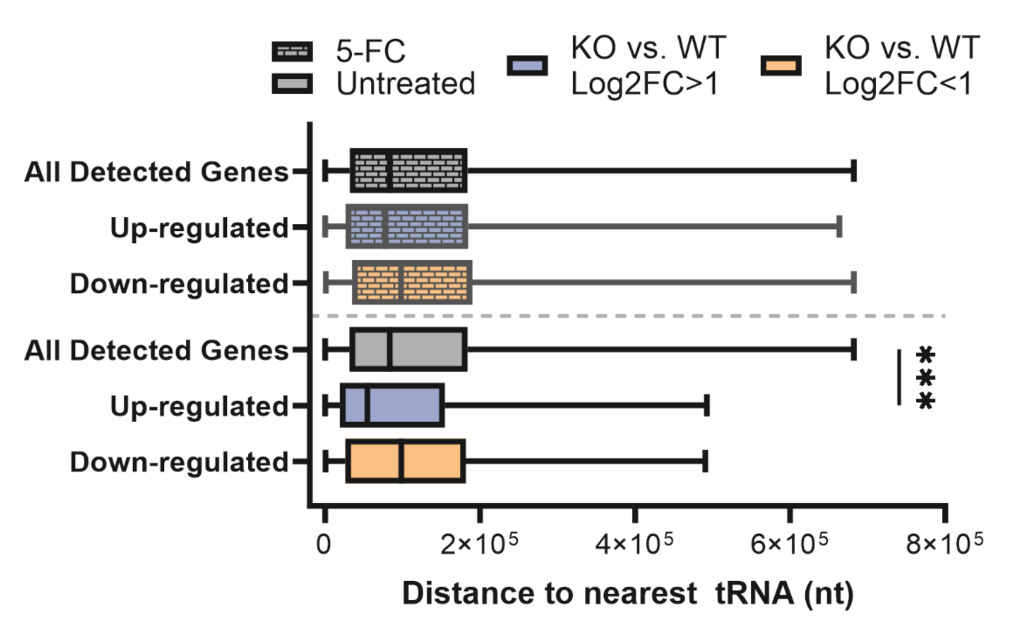


Figure S12. Genes induced in Δ*mod5* display a closer proximity to tRNA genes than down-regulated ones. Transcriptional data of non-stressed (clear boxes) and 5-FC stressed (checked boxes) samples were correlated with the average distances between tRNA genes and protein-coding genes. Significance was determined by Brown-Forsythe and Welch ANOVA test with Dunnett's T3 multiple comparisons test as posttest. P-value ***=0.0004.
